# Microglia-derived TGF-β1 ligand maintains microglia homeostasis via autocrine mechanism and is critical for normal cognitive function in adult mouse brain

**DOI:** 10.1101/2023.07.05.547814

**Authors:** Alicia Bedolla, Elliot Wegman, Max Weed, Aditi Paranjpe, Anastasia Alkhimovitch, Igal Ifergan, Lucas McClain, Yu Luo

## Abstract

While TGF-β signaling is essential for microglial function, the cellular source of TGF-β ligand and its spatial regulation remains unclear in the adult CNS. Our data support that microglia, not astrocytes or neurons, are the primary producers of TGF-β1 ligands needed for microglial homeostasis. Microglia (MG)-*Tgfb1* inducible knockout (iKO) leads to the activation of microglia featuring a dyshomeostatic transcriptomic profile that resembles disease-associated microglia (DAMs), injury-associated microglia, and aged microglia, suggesting that microglial self-produced TGF-β1 ligands are important in the adult CNS. Interestingly, astrocytes in MG-*Tgfb1* iKO mice show a transcriptome profile that closely aligns with A1-like astrocytes. Additionally, using sparse mosaic single-cell microglia iKO of TGF-β1 ligand, we established an autocrine mechanism for TGF-β signaling. Importantly MG-*Tgfb1* iKO mice show cognitive deficits, supporting that precise spatial regulation of TGF-β1 ligand derived from microglia is critical for the maintenance of brain homeostasis and normal cognitive function in the adult brain.

## Introduction

Microglia are commonly known as the resident immune cells in the central nervous system (CNS), but their roles expand beyond that of innate immunity. At homeostasis, microglia play a variety of regulatory roles such as surveilling the brain parenchyma for injury or disease, phagocytosis, and synaptic pruning^1–4^. In addition to their homeostatic role, microglia are vital in inflammatory response initiation and regulation. In the case of injury or inflammation, microglia dynamically alter their function on a spectrum of activation states ranging from the more pro-inflammatory M1-like state to the anti-inflammatory M2-like state^5–7^. Previous studies have shown that transforming growth factor beta (TGF-β) signaling is required for the development of microglia during the embryonic stage^8^. Specifically, a cleverly designed “CNS-specific” *Tgfb1* knockout (KO) mouse model was developed by overexpressing the *Tgfb1* gene in T-cells (via an Il-2 Promoter) in a global *Tgfb1* KO mouse model, which depletes CNS TGF-β1 constitutively but partially compensates peripheral TGF-β1 levels^8^. Using this mouse model, it was reported that in the absence of TGF-β1 in the CNS during development, microglia do not establish their signature gene expression, supporting that TGF-β1 is critical for normal microglial development^8^. While this study supported the importance of TGF-β1 in microglial development, whether TGF-β signaling is required in mature microglia to maintain their survival and function in adult brain is not known. Moreover, the serum levels of TGF-β1 in this “CNS” *Tgfb1* KO mouse model were undetectable^8^, resulting in a potential confound due to altered TGF-β1 levels in peripheral tissues and serum which could have indirect effects on microglia maturation.

Recently, using an inducible myeloid-specific TGF-β type 2 receptor (TGF-βR2) KO mouse model (*Cx3cr1*^CreER^-*Tgfbr2*^fl/fl^), Zoller et al. have shown that TGF-β signaling, via TGF-βR2 in adult microglia, is necessary for maintaining the ramified morphology and certain features of microglial homeostasis^9^. However, inducible knockout (iKO) of *Tgfbr2* in adult microglia only leads to a “primed” state in microglia without any effects on many microglia homeostatic signature genes such as *P2ry12*, *Tmem119*, *Hexb*, and *Sal1*^9^. These studies, while supporting the importance of TGF-β signaling in microglia maturation during developmental stages and maintaining certain features of homeostasis in adulthood, also generate some standing questions regarding the critical requirements of TGF-β signaling in maintaining microglia signature gene expression in the adult CNS. Additionally, the precise cellular source of TGF-β ligands and the spatial/temporal regulation of components of the TGF-β signaling pathway cross different cell types in the CNS and its functional relevance have yet to be identified, leaving some major gaps in our knowledge regarding the regulation of this important signaling pathway in the adult brain.

While it is assumed that many different cell types can be sources of TGF-β ligands in the CNS at homeostasis^10–15^, the actual production of TGF-β ligands in different cell types in the brain has not been established. Additionally, a previous study hints at a highly precise spatially localized regulation of the activation of TGF-β ligand through the interaction of LRRC33 and αVβ8 integrin^16^. However, whether TGF-β ligand is regulated by a diffusible paracrine mechanism locally or whether the ligand is more strictly regulated via an autocrine manner in the CNS is not known. We aim to address these major gaps in the field in this study. Using cell-type specific conditional or inducible KO models of the *Tgfb1* gene, we demonstrate that microglia- but not neuron- or astrocyte-derived TGF-β1 ligand is critical for the maintenance of homeostatic microglia in the adult brain. Furthermore, the loss of microglia-derived TGF-β1 ligands also leads to the presence of reactive astrocytes in the brain and causes cognitive deficits in adult mice. Additionally, our study shows that the total TGF-β1 ligand level is substantially lower in the brain compared to serum or peripheral TGF-β1 levels, and that the adult brain has accordingly established a precise spatially controlled mechanism to regulate ligand production to maintain homeostasis in individual microglia, dependent on microglial autocrine TGF-β signaling. Our data also show that TGF-β1 is enriched in microglia, while TGF-β2 is instead enriched in astrocytes. Interestingly, following *Tgfb1* gene deletion in microglia, *Tgfb3* is upregulated in the *Tgfb1* KO microglia. However, at least up to 8 weeks following global microglial *Tgfb1* deletion, neither the astrocytic *Tgfb2* nor the upregulated *Tgfb3* in microglia is able to compensate for the function of *Tgfb1* or rescue the dyshomeostatic phenotype in *Tgfb1* KO microglia, suggesting distinct expression and functions of the different ligands in CNS. In this study, we also address the questions of microglia-astrocyte crosstalk and the functional relevance of microglial TGF-β signaling in the adult CNS. With the importance of TGF-β signaling becoming more recognized in injury, neurodegeneration, and aging in the CNS, our study provides new insights into the mechanisms of how TGF-β signaling can be regulated on a single-cell level via microglia autocrine mechanism in the adult CNS and opens new directions for future studies in understanding how TGF-β1 ligand production and downstream signaling in recipient cells can occur under these conditions.

## RESULTS

### Myeloid-specific Tgfb1 gene deletion in CNS microglia leads to decreased microglial TGF-β signaling without affecting spleen or serum TGF-β1 level in the Cx3cr1^CreER^-Tgfb1 iKO mouse line

To identify the cell type(s) in the CNS that provide TGF-β1 ligand to microglia and other TGF-β1 responsive cells, we first examined scRNAseq data sets published in previous studies^17,18^. Interestingly, highly enriched *Tgfb1* mRNA levels in adult mouse microglia are observed in multiple scRNAseq datasets^17,18^. Further analysis of these data shows that astrocytes, neurons, and oligodendrocytes have minimal *Tgfb1* expression. While OPCs and endothelial cells have moderate *Tgfb1* expression, it is substantially lower compared to microglia *Tgfb1* mRNA levels (Supplementary Fig 1A,B)^17,18^. To validate microglia *Tgfb1* expression, we carried out a combined RNAscope/IHC analysis to examine the cellular expression pattern of TGF-β1 in the adult mouse brain. Our data show that indeed *Tgfb1* mRNA is enriched in IBA1 positive microglia, but was not detected in neurons (Supplementary Fig 1C). *Tgfb1* mRNA is also detected in a small population of non-IBA1+ cells, which could be endothelial cells or OPCs. Interestingly, mRNA for the type 1 TGF-β1 receptor (TGF-βR1 or ALK5) is also detected in microglia, supporting that microglia can be both ligand-producing and responding cells for TGF-β1 signaling. To examine whether microglia are a major contributor to TGF-β1 production in the brain, we depleted microglia from the adult mouse brain using the well-established CSFR1 antagonist PLX5622 (1200ppm in diet) following our previously published protocol, which resulted in more than 90% of microglia ablation reported by us and other studies^19–21^. After successful microglia ablation, we examined the total *Tgfb1* mRNA in cortical tissue from control or microglia-ablated mouse brain. The qRT-PCR analysis shows that the PLX5622 treatment leads to a substantial depletion of microglia in the adult brain, indicated by a significant decrease of *Iba1* mRNA levels in brain tissue (<5% of WT levels), accompanied by a 70% decrease in total *Tgfb1* mRNA levels (Supplementary Fig 1E). Since microglia compose only 5-10% of total brain cells^22,23^, microglia ablation leading to a 70% decrease in total *Tgfb1* mRNA levels in the brain supports that microglia are a major component for TGF-β1 ligand production.

Next, to establish direct functional relevance of microglia-produced TGF-β1 ligand to microglia homeostasis, the *Cx3cr1*^CreER^ line^24^ was crossed with the *Tgfb1*^fl/fl^ line to enable tamoxifen (TAM)-induced TGF-β1 ligand loss in microglia in adulthood. To confirm the efficiency of *Tgfb1* gene deletion in microglia in the inducible MG-*Tgfb1* KO mice, in our recent study, we sorted microglia using an R26-YFP reporter allele (which labels 90% of total microglia in adult mouse brain) at 3 weeks following TAM treatment in *Cx3cr1*^CreER^*Tgfb1^wt/wt^R26-YFP and Cx3cr1*^CreER^*Tgfb1^fl/fl^R26-YFP* mice^25^. We demonstrate a significant decrease (99.8%) in *Tgfb1* mRNA levels from sorted YFP+ microglia in *Cx3cr1*^CreER+/-^-*Tgfb1*^fl/fl^ mice in comparison to control microglia (*Cx3cr1*^CreER+/-^, *Tgfb1*^wt/wt^)^25^. To examine whether peripheral serum or tissue TGF-β1 levels are also affected in our MG-*Tgfb1* iKO mice, we measured the TGF-β1 protein levels in the serum and spleen using ELISA analysis. In contrast to the previous “CNS-*Tgfb1*” constitutive KO mouse model which completely abolished serum TGF-β1 levels^8^, our *Cx3cr1*^CreER^*Tgfb1* iKO mice do not show any difference in TGF-β1 protein levels in the serum or spleen between the Cx3cr1^CreER^ WT and the *Cx3cr1*^CreER^*Tgfb1*^fl/fl^ mice at 3 weeks after TAM treatment (Supplemental Fig 2), confirming the minimal interference in peripheral TGF-β1 ligand levels. Interestingly, ELISA results also show that compared to the high levels of TGF-β1 ligand in the spleen and serum (Control and iKO Spleen = 5.7 or 5.9pg/ug protein: Control and iKO serum= 1.9 or 1.8pg/ug protein), total brain TGF-β1 levels are substantially lower (below the detect limit of the ELISA assay), a result that is consistent with a recent previous study^25^. Due to the limitation of this analysis method, we were not able to confirm loss of TGF-β1 ligand on a protein level in our *Cx3cr1-Tgfb1* iKO mice directly using ELISA assay. Similarly, our results were also confirmed using FACS analysis on surface TGF-β1 levels. FACS analysis showed no difference in TGF-β1 expression levels in CD11b+/CD45+ spleenocytes between control and *Cx3cr1^CreER^Tgfb1* iKO mice, while CD11b+/CD45+ cells from the brain show very low levels of TGF-β1 signal. This further supports that CNS TGF-β1 levels are substantially lower compared to peripheral tissue, making detection of the TGF-β1 protein in the CNS a challenge. Due to the very low signal of TGF-β1 staining in brain CD11b+/CD45+ cells, we could not compare the level of surface TGF-β1 expression between control and *Cx3cr1^CreER^-Tgfb1* iKO mice. Nevertheless, the qRT-PCR analysis from sorted brain microglia demonstrates that the *Cx3cr1*^CreER^*Tgfb1*^fl/fl^ transgenic line can efficiently delete the *Tgfb1* gene in microglia cells specifically with minimal interference on systemic serum or spleen TGF-β1 levels^25^. Additionally, RNAseq of the sorted WT and *Tgfb1* iKO microglia confirms the complete loss of the floxed exon 3 from *Tgfb1* mRNA in iKO mice while not affecting mRNA counts of exon 4 which is downstream of the 3’ loxP site (Fig 1 E).

**Figure 1.**
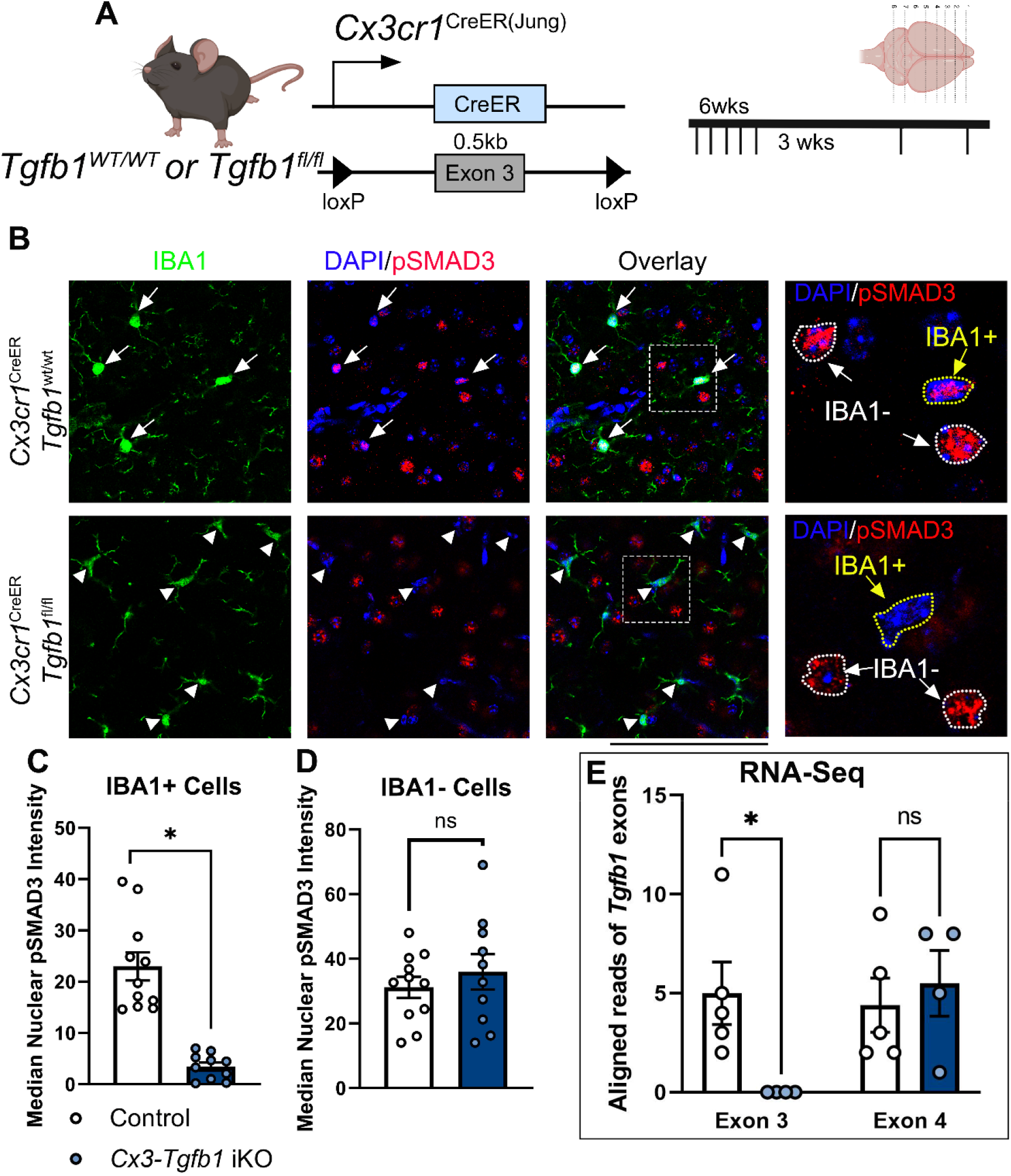
Validation of loss of *Tgfb1* loxP-flanked exon 3 in microglial mRNA and the TGF-β downstream signaling effector (pSMAD3) in *Cx3cr1*^CreER(Jung)^*Tgfb1*^fl/fl^ iKO mice following tamoxifen administration. (A) Mouse model used and experimental timeline. (B) Representative IHC showing pSMAD3, IBA1, and DAPI in control animals and iKO animals. (C,D) Quantification of pSMAD3 nuclear immunoreactivity median intensity in both IBA1+ cells (C) and IBA1-cells (D). (E) Based on RNAseq analysis of sorted MG from Control or *Cx3cr1*^CreER(Jung)^*Tgfb1*^fl/fl^ iKO mice, aligned reads that matched to exon3 (the loxP-flanked exon) or exon4 (the exon downstream of the floxed exon) show that exon 3 is significantly lower in MG-*Tgfb1* iKO microglia compared to control microglia while exon 4 is unaffected. Mean±SE, * = p<0.05. Student’s t-test. Scale bar = 100µm. SMAD3 bar graphs show individual image averages, however statistics were carried out using the average cell intensity for a single animal (control n=4, iKO n=3). RNA-seq graph shows a single data point per animal.

To further circumvent the challenge of direct detection of TGF-β1 protein, we analyzed the downstream effector of TGF-β1 signaling, i.e., the nuclear localized phosphorylated SMAD3 (pSMAD3) protein levels. Co-immunohistochemistry analysis shows that in control mice (*Cx3cr1*^CreER+/-^*Tgfb1*^wt/wt^ + TAM), pSMAD3 is detected in both IBA1+ microglia cells and IBA1-cells in the brain (Fig 1B, C, D). Interestingly, MG-specific deletion of the *Tgfb1* gene leads to a specific and significant decrease of pSMAD3 immunoreactivity exclusively in IBA1+ microglia, without affecting the pSMAD3 immunostaining in IBA1-cells (Fig 1 B-D). This specific loss of TGF-β1 signaling (pSMAD3) in microglia from MG-*Tgfb1* iKO mice confirms that microglia TGF-β1 signaling depends on microglia produced TGF-β1 ligand which cannot be compensated for by other cells.

### Microglia-derived TGF-β1 ligand is critical to maintain microglial homeostasis and astrocyte quiescence in the adult CNS

Next, we evaluated whether the loss of microglia-derived TGF-β1 ligand affects microglia morphology and homeostatic status. At 5- and 8-weeks post TAM administration, substantial morphological changes in microglia in MG-*Tgfb1* iKO mice were observed (Fig 1B and Fig 2D, E - IBA1 staining in green). This change in morphology of IBA1+ cells in the iKO mice suggests loss of ramification and potential activation of microglia, which prompted us to carry out further detailed morphological analysis and examination of the homeostatic microglia signature genes. Two independent Cx3cr1^CreER^ mouse drivers were used in this study to confirm the phenotypes. Following microglial *Tgfb1* knockout, microglia in *Cx3cr1*^CreER(Littman)24^ *Tgfb1* iKO mice showed less ramification indicated by decreased total branch length (Fig 3F, control=323µm, 5wk iKO=167µm, 8wk iKO=190µm) and processes terminal end number (control=26, 5wk iKO=16, 8wk iKO=17) compared to control microglia. Moreover, *Tgfb1* iKO microglia showed decreased expression of homeostatic microglia signature genes such as *P2ry12* and *Tmem119* (Fig 2C-E) and an upregulation of CD68 expression. These results agree with an impaired microglial homeostatic status; however, our observed phenotype is more severe than that of a previous study in which adult microglial TGF-β signaling was abolished via KO of TGF-βR2^9^, supporting that loss of microglia-derived TGF-β1 ligand cannot be compensated by potential TGF-β1 ligand production in other cell types in the adult CNS. These results support that microglial TGF-β1 signaling relies on TGF-β1 ligand produced by microglia. Interestingly, an increase in reactive astrocytes (indicated by upregulated GFAP expression, Fig 2I) was also observed both at 5 weeks and 8 weeks after TAM treatment in the *Cx3cr1*^CreER(Littman)^ *Tgfb1*^fl/fl^ mice but not in the *Cx3Cr1*^CreER(Littman)^*Tgfb1*^wt/wt^ +TAM or the cre(-)*Tgfb1*^fl/fl^ +TAM mice. We first examined and quantified the phenotype in microglia and astrocytes in the cortex of the *Cx3cr1*^CreER(Littman)^*Tgfb1*^fl/fl^ line (Fig 2) and observed a similar phenotype for both microglia and astrocytes at other brain regions as well (Supplementary Fig 3 showing hippocampus as another example, note that unlike cortical astrocytes which are mostly GFAP-, hippocampal astrocytes are already GFAP+ during homeostasis in WT mice).

**Figure 2.**
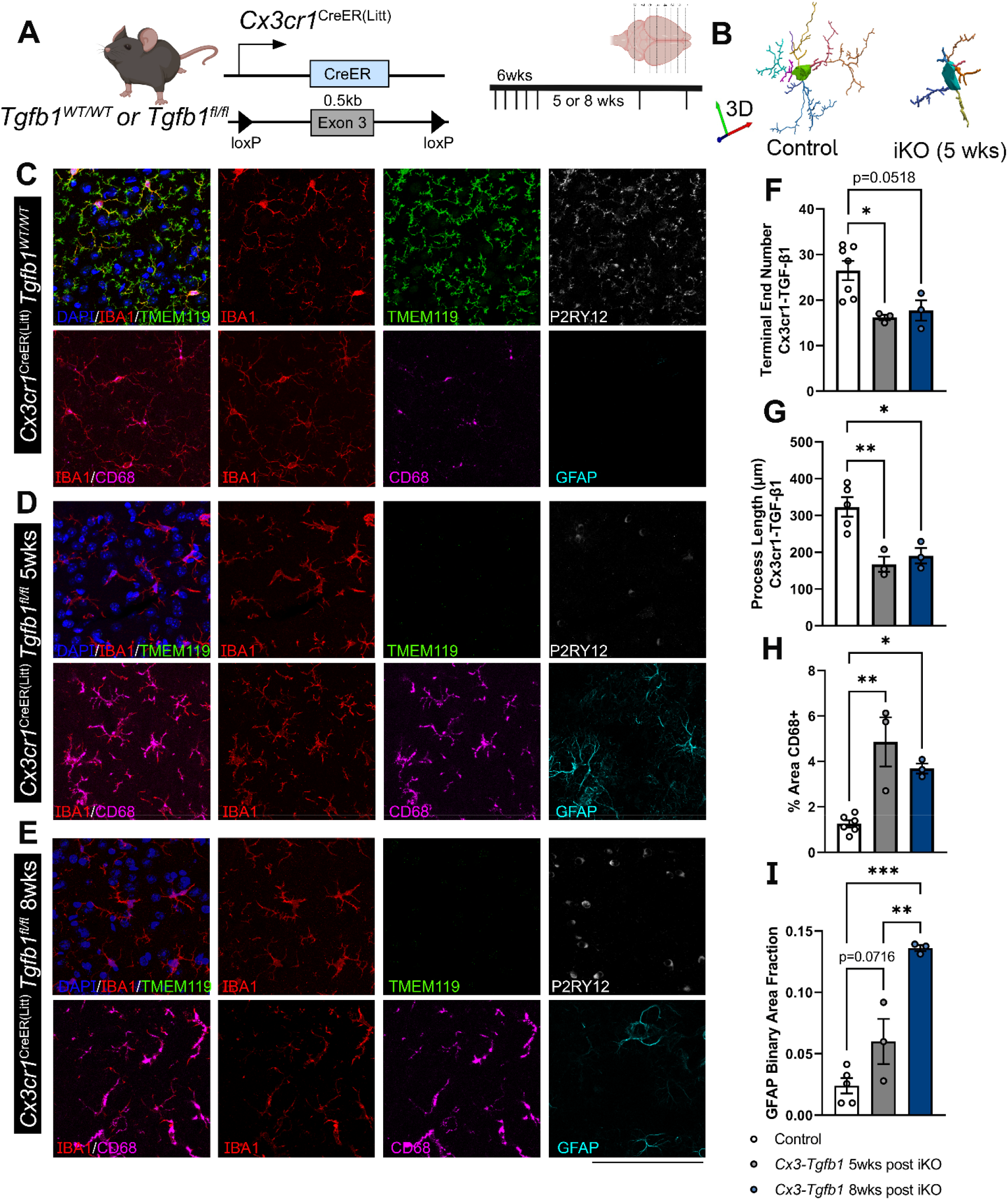
Microglia-specific *Tgfb1* gene deletion results in loss of homeostasis of microglia and in increased reactive astrocytes in cortex of the adult mouse brain. (A) Mouse model for targeting microglial *Tgfb1* and timeline. (B) 3D reconstruction of control and iKO microglia. Representative immunohistochemistry images of IBA1, TMEM119, P2RY12, CD68, and GFAP in the cortex of (C) Control animals, (D) *Cx3cr1*^CreER(Litt)^*Tgfb1*^fl/fl^ knockouts 5 weeks after tamoxifen administration, and (E) *Cx3cr1*^CreER^ ^(Litt)^*Tgfb1*^fl/fl^ knockouts 8 weeks after tamoxifen administration. Quantification of (F) total microglial process length, (G) microglial process terminal end numbers, (H) % of CD68 immunoreactive positive area, and (I) GFAP immunoreactive positive area fraction. Mean±SE, * = p<0.05, ** = p<0.01, *** = P<0.001. Student’s t-test. (> 40 microglia were quantified for each animal and the average from one mouse was plotted as a single data point in the figure panel and treated as n=1 for statistical analysis). Scale bar = 100µm.

**Figure 3.**
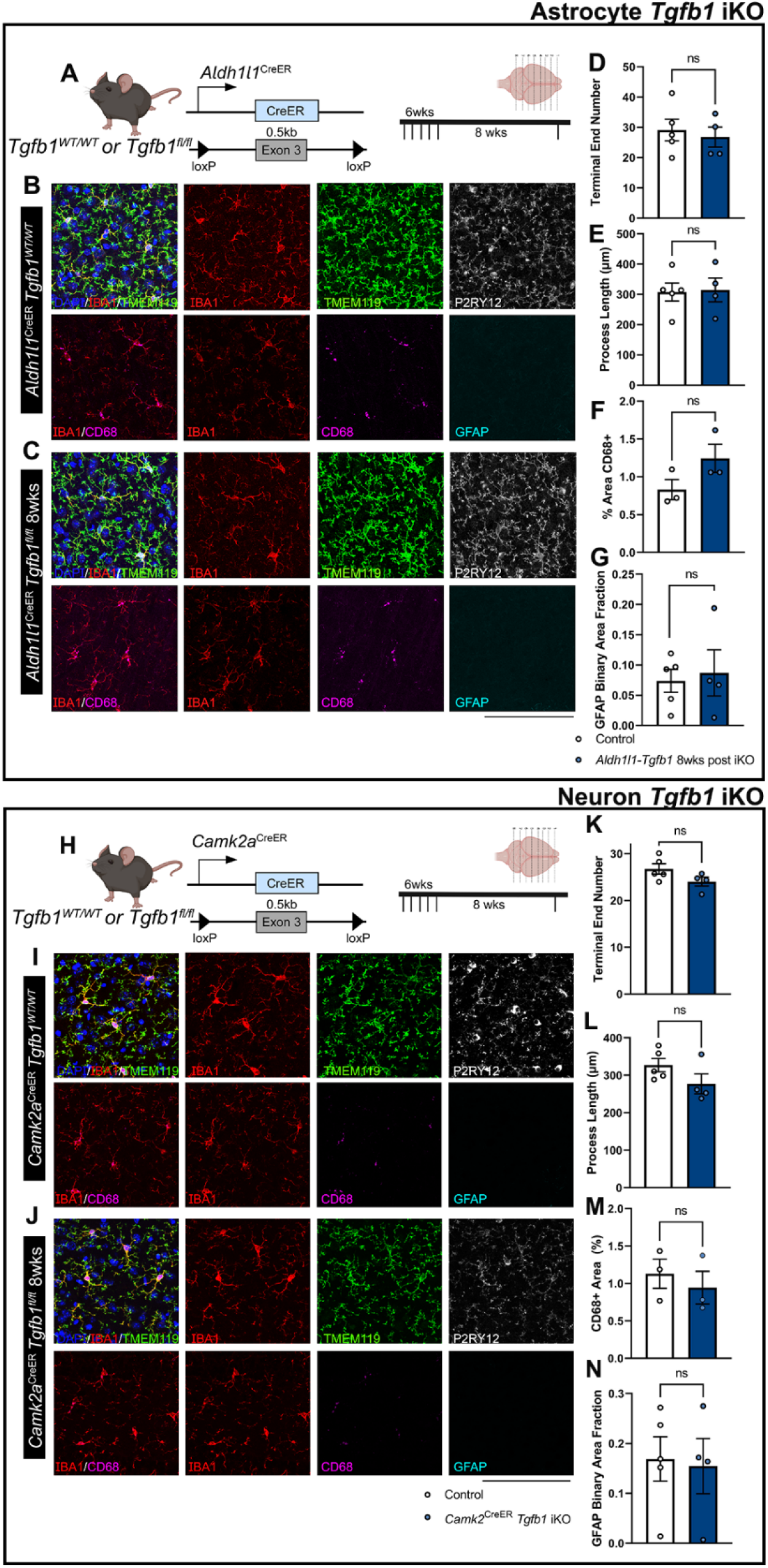
Astrocyte-specific or forebrain neuronal specific *Tgfb1* gene deletion in the *Aldh1l1*^CreER^ or *Camk2a*^CreER^ drivers does not affect the homeostasis of microglia or GFAP expression in astrocytes in adult mouse brain (cortex). (A) Astrocyte iKO mouse model and experimental timeline. (B, C) Representative immunohistochemistry images of cortex from TAM treated (8 weeks post) control (B) *Aldh1l1*^CreER^*Tgfb1*^wt/wt^ and (C) iKO *Aldh1l1*^CreER^ *Tgfb1*^fl/fl^ tissue showing IBA1, TMEM119, P2RY12, CD68, and GFAP immunostaining. Quantification of microglia ramification via (D) process terminal end numbers, (E) total process length, and (H) % of CD68+ immunoreactive area. (G) quantification of astrocyte reactivity using GFAP immunoreactive positive area fraction. (H) Neuronal iKO mouse model and experimental timeline. (I, J) Representative images of TAM treated (8 Weeks post) control *Camk2a*^CreER^*Tgfb1*^wt/wt^ (I) and iKO *Camk2*^CreER^ *Tgfb1*^fl/fl^ tissue showing IBA1, TMEM119, P2RY12, CD68, and GFAP immunoreactivity. Quantification of microglia ramification via (K) process terminal end number, (L) total process length, and (M) CD68+ immunoreactive % area. (N) Quantification of astrocyte reactivity using GFAP+ immunoreactive area fraction. Mean±SE (> 40 microglia were quantified for each animal and the average from one mouse was plotted as a single data point in the figure panel and treated as n=1 for statistical analysis). ns=not significant., Scale bar = 100µm.

During the course of this study, we observed and reported non-TAM dependent leakiness in floxed reporter alleles in the *Cx3cr1*^CreER(Littman)^ line^24^ and became aware of TAM-induced off-target cre activity in this line caused by neonatal TAM injections which could potentially confound our observed microglial phenotype^26^. Although we did not treat control or *Cx3cr1*^CreER(Littman)^*Tgfb1* iKO mice until the age of 6-8 weeks, to further increase the rigor of our study, we generated a second independent MG-*Tgfb1* iKO line using the *Cx3cr1*^CreER(Jung)^ line^27^, in which we and others have recently demonstrated less non-TAM dependent leakiness in reporter alleles and the absence of TAM-induced neonatal or adult off-target effects^25,28^. Using this second independent *Cx3cr1*^CreER^*Tgfb1* iKO mouse line, we confirmed the same microglia and astrocytes activation phenotype in *Cx3cr1*^CreER(Jung)^*Tgfb1*^fl/fl^ mice treated with TAM, while two control groups i.e. *Cx3cr1*^CreER(Jung)^*Tgfb1*^wt/wt^ + TAM treatment or *Cx3cr1*^CreER^*Tgfb1*^fl/fl^ + vehicle injections show no phenotype (Supplemental Fig. 4). All subsequent experiments for microglia-specific *Tgfb1* iKO were therefore carried out in the *Cx3cr1*^CreER(Jung)^*Tgfb1*^fl/fl^ mouse line.

### TGF-β1 ligand produced by astrocytes is not necessary for maintenance of glial homeostasis

Previous studies have also suggested that TGF-β ligands produced by astrocytes could also play important roles in blood brain barrier formation, stabilization, and maturation, as well as neuroprotection following injury or disease^29–31^. Therefore, we next investigated whether astrocyte-specific deletion of the *Tgfb1* gene would also lead to alterations in microglia morphology and gene expression changes such as *P2ry12*, *Tmem119*, and *Cd68*. Additionally, we aimed to investigate whether loss of the astrocytic *Tgfb1* gene could also induce changes in astrocyte reactivity as observed in *Cx3cr1*^CreER^*Tgfb1* iKO mice. To target adult astrocytes, *Aldh1l1*^CreER^ mice^32^ were crossed with *Tgfb1*^fl/fl^ mice to generate astrocytic *Tgfb1* iKO mice (*Aldh1l1*^CreER^*Tgfb1*^fl/fl^). At 8-weeks following TAM administration, microglia morphology and astrocyte state were analyzed. Microglial morphology remained unchanged in *Aldh1l1*^CreER^*Tgfb1*^fl/fl^ animals compared to control mice (*Aldh1l1*^CreER^*Tgfb1*^wt/wt^ mice or *Tgfb1*^fl/fl^ mice) (Fig 3 for cortex and Supplementary Fig 5 for hippocampus as example). Additionally, no changes were observed in homeostatic microglia signature genes (*P2ry12* or *Tmem119*). Nor did we observe upregulation of CD68 in microglia or an increase in GFAP expression in astrocytes when comparing the *Aldh1l1*^CreER^*Tgfb1*^fl/fl^ animals to wildtype controls at 8 weeks after TAM treatment (Fig 3 F, G and Supplementary Fig 5). This result further confirms that astrocytic TGF-β1 production is not required for the maintenance of microglia and astrocyte homeostasis.

Additionally, we also generated the constitutive astrocytic *Tgfb1* KO mice using the *mGfap*^cre^ driver line^33^, which targets a large population of astrocytes constitutively starting from neonatal stages^33^, to ensure that a larger population of astrocytes (95% of cortical astrocytes labeled with Ai14 reporter)^34^ will have the TGF-β1 ligand KO. Comparing the morphology of *mGfap*^Cre^*Tgfb1*^fl/fl^ microglia to wildtype controls, there were no changes in the ramification of microglia in this independent astrocytic-*Tgfb1* cKO mouse line (Supplementary Fig 6 for cortex and Supplementary Fig 7 for hippocampus). We next examined the expression of the homeostatic microglia signature genes such as *P2ry12* and *Tmem119* and did not observe any difference between *mGfap*^cre^*Tgfb1*^fl/fl^ mice and control mice. CD68 expression in microglia and astrocytic GFAP expression were also unchanged. These results together support that under normal physiological conditions, adult astrocytes do not produce TGF-β1 ligand necessary for homeostatic maintenance in both microglia and astrocytes. It remains to be determined whether under injury or pathological conditions astrocytes could upregulate TGF-β1 ligand to modulate glial responses to injury or neurodegeneration.

### Forebrain excitatory neurons do not produce TGF-β1 ligand for neighboring microglia

Next, we investigated whether neurons are an additional critical source for *TGF-β1* ligand production for adult microglia. To this end, we generated a forebrain excitatory neuron-specific *Tgfb1* iKO mouse model. To target forebrain neurons, a *Camk2a*^CreER^ line^35^ was crossed with the *Tgfb1*^fl/fl^ line to induce TGF-β1 ligand KO in excitatory neurons^35^. Camk2a^CreER^ has been reported to recombine in a widespread manner in the cortex, hippocampus, and striatum^35^. Eight weeks after TAM administration, microglial morphology remained unchanged in the *CamkII*^CreER^*Tgfb1*^fl/fl^ mice compared to wildtype controls (Fig 3 for cortex and Supplementary Fig 8 for hippocampus). Additionally, no alterations in TMEM119 or P2RY12 expression in microglia nor an increase in GFAP expression was observed. This supports that microglia do not critically rely on neuronal TGF-β1 ligand to maintain homeostasis in the adult brain.

### Tmem119^CreER^ and P2ry12^CreER^ microglia-specific CreER mouse lines lead to lower-efficiency gene deletion of TGF-β1 in subsets of microglia and result in mosaic activation of adult brain microglia

The *Cx3cr1*^CreER^ line has previously been reported to also target border associated macrophages (BAMs) that reside in the pia and vasculature^25,28^. Additionally, the two *Cx3cr1*^CreER^ mouse lines replace the endogenous *Cx3cr1* gene with the cre expression cassette, resulting in heterozygosity for the *Cx3cr1* gene in both control (*Cx3cr1*^CreER^*Tgfb1*^wt/wt^) and in iKO (*Cx3cr1*^CreER^*Tgfb1*^fl/fl^) mice^24,27^. In an effort to further improve the targeting specificity for parenchymal microglia, the *Tmem119*^CreER^ and *P2ry12*^CreER^ mouse lines have recently been generated, utilizing homeostatic microglia signature gene promoters to drive creER cassette expression without affect the endogenous gene expression of *Tmem119* or *P2ry12*^36,37^. Our recent study reports that the *Tmem119*^CreER^ and the *P2ry12*^CreER^ mouse lines do show less TAM-independent “leaky” recombination events with the drawback of lower recombination efficiency, leading to only a subset of microglia being recombined in these two mouse lines (based on R26-YFP reporter gene expression and qRT-PCR results in sorted microglia)^25,28^. To examine whether mosaic gene deletion of the TGF-β1 ligand in a subset of parenchyma microglia would generate any phenotype in microglia, we crossed *Tmem119*^CreER^ and *P2ry12*^CreER^ with the *Tgfb1*^fl/fl^ line to generate two parenchymal-microglia specific *Tgfb1* iKO lines.

Our recent study shows that the R26-YFP reporter (which has a larger floxed stop cassette between the two loxP site, 2.7kb) has more stringent TAM-regulated cre recombination compared to the Ai9 R26-CAG-tdTomato reporter (0.9 kb between two loxP sites). Our study also shows that compared to the *Cx3Cr1*^CreER^*Tgfb1* iKO line, *P2ry12*^CreER^*Tgfb1* iKO mice show significantly lower efficiency of *Tgfb1* gene deletion (∼50% decrease in floxed *Tgfb1* exon 3 mRNA levels) in sorted YFP+ microglia cells after TAM treatment^25^. To investigate whether this incomplete deletion of microglial *Tgfb1* could still produce the disruption of microglial homeostasis we observed in the *Cx3Cr1*^CreER^*Tgfb1* iKO lines, we next evaluated microglial morphology in *P2ry12*^CreER^ and *Tmem119*^CreER^*Tgfb1* iKO mice. Consistent with the mosaic recombination of the R26-YFP reporter gene and the partial reduction of *Tgfb1* mRNA levels in adult microglia in the *P2ry12*^CreER^ mouse line, three weeks after TAM administration, we observed distinct islands of YFP+ microglia patches with activated microglia morphology in both the *Tmem119*^CreER^*Tgfb1* and *P2ry12*^CreER^*Tgfb1* iKO mice (Fig 4B, D, E). In addition to the mosaic and patchy morphological changes, we also observed a decrease in expression of homeostatic microglia signature genes *P2ry12* and *Tmem119* in these patches of microglia (Fig 4B, D, E). The morphological changes and loss of *P2ry12* and *Tmem119* expression were not observed in control *Tmem119*^CreER^*Tgfb1^wt/wt^*or *P2ry12*^CreER^*Tgfb1^wt/wt^* TAM mice or in *Tmem119*^CreER^*Tgfb1^fl/fl^* and *P2ry12*^CreER^*Tgfb1^fl/fl^* mice treated with vehicle only. Notably, the distinct mosaic patches of the few *Tgfb1* KO microglia which showed altered morphology and loss of expression of homeostatic microglia signature genes are surrounded by wildtype microglia cells which can produce the TGF-β1 ligand (Fig 4B, D, E). This raises the interesting question of whether individual microglia rely on self-produced TGF-β1 ligand that is secreted in an autocrine manner or whether individual microglia could utilize the TGF-β1 ligand from neighboring microglia. To further investigate the mechanism of microglia-produced TGF-β1 ligand via an autocrine or paracrine mechanism, we further designed a sparse labeling strategy using the more robust *Cx3cr1*^CreER^ mouse line with a diluted TAM regimen as detailed below.

**Figure 4.**
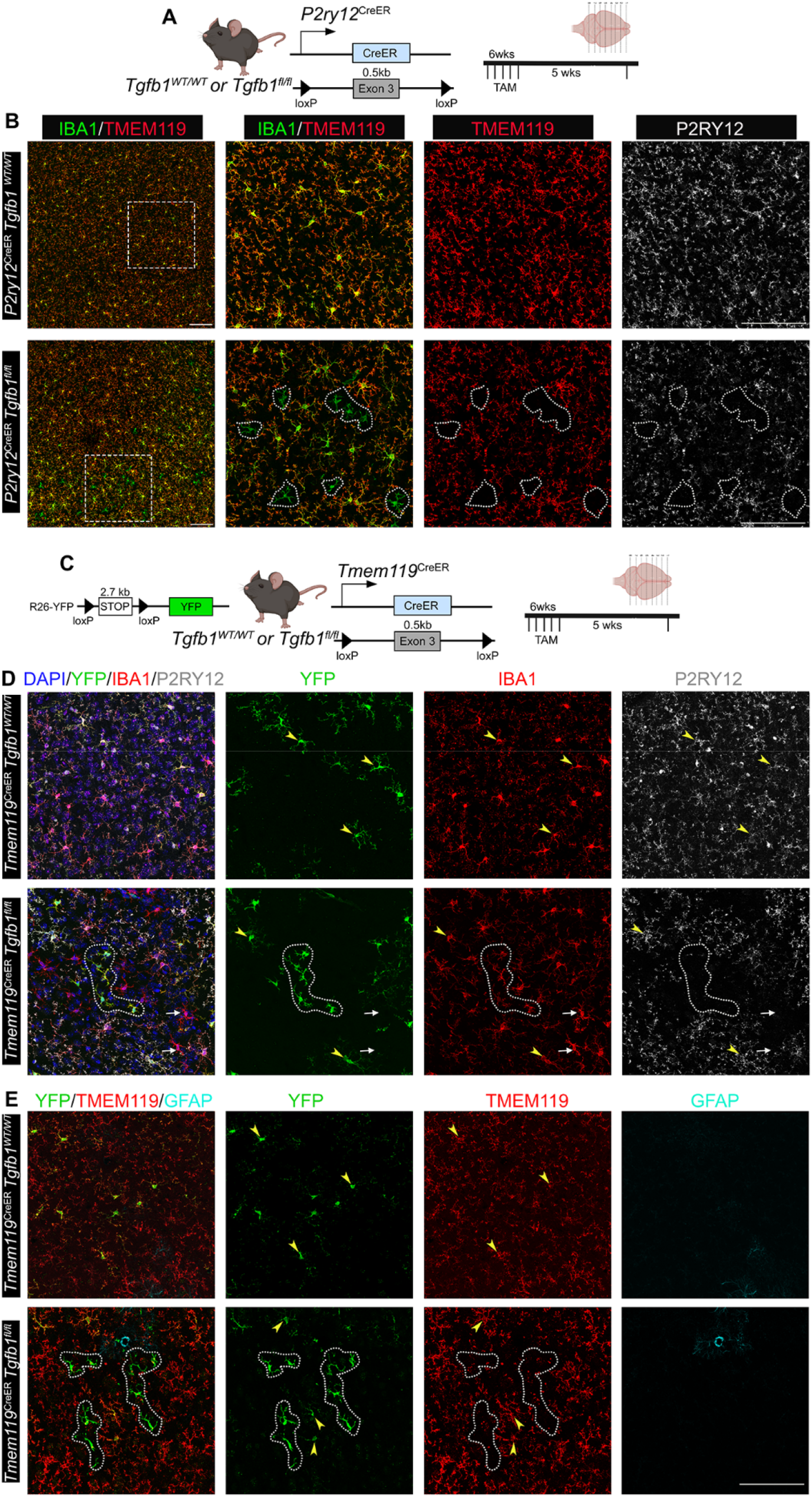
Mosaic deletion of *Tgfb1* gene in subsets of parenchyma microglia in the *P2ry12*^creER^ *Tgfb1* ^fl/fl^ or the *Tmem119*^CreER^*Tgfb1*^fl/fl^ iKO mice leads to distinct patches of dyshomeostatic microglia in the adult mouse brain. (A) P2ry12^CreER^ mouse driver to induce *Tgfb1* KO in P2RY12+ microglia and experimental timeline. We recently showed that these lines have a 50% decrease in exon3 of *Tgfb1* mRNA in reporter positive microglia^25^. (B) TAM treated (5 weeks post) Control and *P2ry12*^CreER^*Tgfb1* ^fl/fl^ iKO representative images showing immunohistochemistry for IBA1, TMEM119, and P2RY12. (C) Tmem119^CreER^ mouse driver to induce tgfb1 KO in TMEM119+ microglia and experimental timeline. (D) TAM treated (5 weeks post) Control and *Tmem119*^creER^*Tgfb1* ^fl/fl^ iKO representative images showing immunohistochemistry for YFP, IBA1, P2RY12 and (E) YFP, TMEM119, and GFAP immunostaining. White dotted outlines indicate microglia with downregulated TMEM119 and/or P2RY12 expression. White arrows depict YFP-cells that also show loss of P2RY12 expression. Yellow arrowheads show YFP+ cells in either WT or iKO mice that still maintained P2RY12 or TMEM119 expression. Representative results from n=3-5 mice/group. Scale bar = 100µm.

### Mosaic knockout of TGF-β1 ligand in sparse single microglia reveals the autocrine mechanism of TGF-β1 signaling regulation that provides precise spatial and temporal regulation of microglia homeostasis

To further investigate the spatial resolution of TGF-β1 ligand production by individual microglia and investigate whether individual microglia rely on self-produced TGF-β1 ligand using an autocrine mechanism, we designed a mosaic sparse recombination strategy using the *Cx3cr1*^CreER(Jung)^*Tgfb1*^fl/fl^ line with a titrated TAM dilution. To accomplish this, we tested a TAM dosage of 1:50 and 1:7-1:10 of the concentration (180mg/kg) that is utilized in our full dose recombination. Our results (Supplementary Fig 9) show that for both the 1:50 and 1:7-1:10 TAM dosages, we observed very sparse YFP+ cells in the parenchyma that are also P2RY12+ (suggesting sparse recombination can occur in parenchymal microglia instead of BAMs). This result supports the feasibility of inducing sparse gene deletion in individual microglia surrounded by WT microglia using a titration of TAM dosage. Since the 1:7-1:10 dosage range provided recombination events that are sufficiently sparse, we carried out our subsequent experiments with this range of dosage. A diluted dose of 1:7 (18mg/kg) of TAM was given over the course of 3 days to *Cx3cr1*^CreER(Jung)^*Tgfb1*^fl/fl^R26-YFP or control mice. This resulted in sparse labeling of microglia in the *Cx3cr1*^CreER(Jung)^ line, allowing for single-cell analysis of whether microglia depend on self-secreted TGF-β1 ligand to maintain their homeostasis. Remarkably, at 2-3 weeks post TAM administration, we observed isolated sparse individual IBA1+ cells in the parenchyma of the *Cx3cr1*^CreER(Jung)^*Tgfb1*^fl/fl^ TAM-treated mice that presented with altered morphology (less ramified), accompanied by a decrease in TMEM119 and P2RY12 expression (Fig 5C, supp Video 1). Interestingly, these sparse mutated single IBA1+ cells are in the brain parenchyma and do not show typical blood vessel-associated macrophage morphology (Fig 5C). Some of these single morphologically-changed, TMEM119-microglia are YFP+ but they are not exclusively YFP+ suggesting that at this low level of TAM dosage, the recombination of the R26-YFP allele can happen independently from the deletion of targeted floxed genes on a single cell level, a result that is also supported by our recent study using dual reporter alleles in microglia^25^. In *Cx3cr1*^CreER^*Tgfb1*^wt/wt^R26-YFP TAM-treated mice, we observed a similar frequency of sparse YFP+ microglia but did not find any individual non-BAM microglia that show this phenotype of activated microglia (marked by loss of TMEM119 expression and altered morphology), suggesting that the sparse individual “activated” microglia in the *Cx3cr1*^CreER^*Tgfb1*^fl/fl^ mice is due to loss of TGF-β1 ligand in these single sparse microglia.

**Figure 5.**
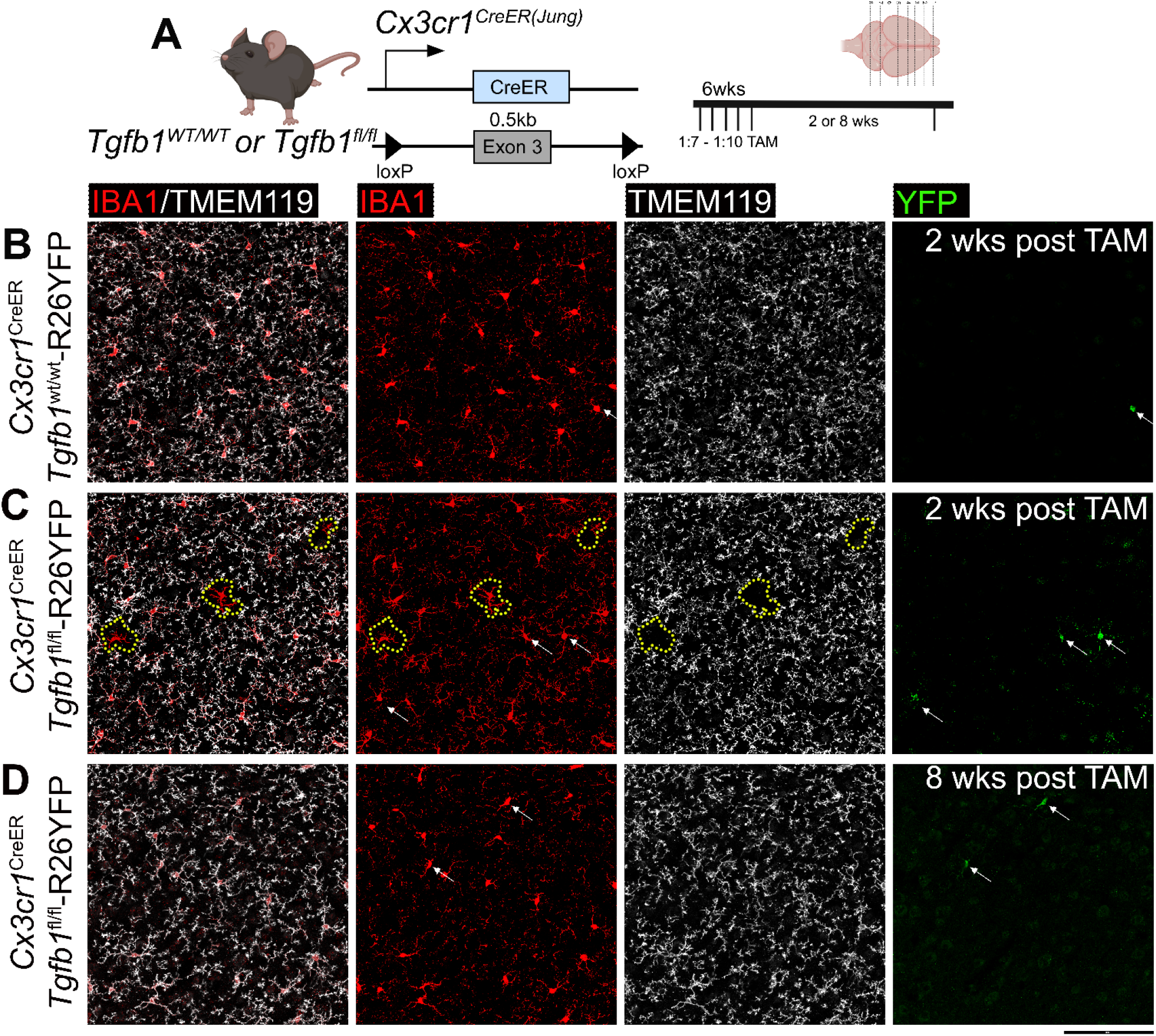
Sparse *Tgfb1* induced-knockout in individual adult microglia supports an autocrine mechanism of microglial TGF-β ligand production and signaling regulation. (A) Mouse model used to induce *Tgfb1* KO in mosaic sparse individual microglia and experimental timeline depicting titrated dose of tamoxifen. (B-D) Representative images showing IBA1, TMEM119, and YFP expression and co-localization in control tissue at 2 weeks post tamoxifen (B) and sparse iKO tissue at 2 weeks (C) showing loss of TMEM119 expression in sparse individual microglia and (D) reversal of TMEM119 expression in the sparse *Tgfb1* iKO brain at 8 weeks post tamoxifen. The yellow dotted outline in (C) highlights singular microglia showing loss of homeostatic TMEM119 expression. White arrows highlight YFP+ cells showing no loss of homeostatic TMEM119 expression. Note that at this low dosage of TAM, the recombination of individual floxed alleles (R26-YFP reporter or the floxed *Tgfb1* gene) occurs independently of each other, therefore YFP+ cells do not always indicate a sparse *Tgfb1* KO microglia, consistent with our recent study^25^. Representative results from n=3-5 mice/group from different cohorts of TAM treatment. Scale bar = 100µm.

To further confirm this hypothesis, we carried out a combined IHC/RNAscope using the same TGF-β1 RNAscope probe we used to confirm microglial TGF-β1 expression (Fig 6B). Indeed, we observed loss of TGF-β1 RNAscope probe hybridization in the individual microglia that specifically showed altered morphology accompanied by the loss of TMEM119 expression, while surrounding IBA1+/TMEM119+ microglia showed normal TGF-β1 RNAscope signal (Fig 6B). We further carried out immunostaining for the detection of the downstream signaling effector of TGF-β1 signaling (pSMAD3) in the mosaic sparse MG-*Tgfb1* iKO brain and confirmed that loss of pSMAD3 is detected specifically in sparse individual microglia that are TMEM119- and morphologically altered (Fig 6C and Supplementary Fig10-11 for additional image examples). Therefore, our results suggest that microglia show a precise spatial regulation of autocrine TGF-β1 signaling reliant on self-produced TGF-β1 ligands under homeostatic physiological conditions. We next asked whether this loss of homeostasis in individual *Tgfb1* KO microglia at 2-3 weeks post TAM is sustained with time or whether, with surrounding wildtype microglia, the individual *Tgfb1* KO microglia can regain homeostasis. Interestingly, at 8 weeks post TAM treatment in sparse MG-*Tgfb1* iKO mice, the phenotype of sparse TMEM119 negative and morphologically altered microglia are no longer observed (Fig 5D). This suggests that these sparse *Tgfb1* KO microglia can regain homeostasis (indicated by normal ramified morphology and restoration of TMEM119 expression) at 8 weeks after TAM treatment in a sparse mosaic MG-*Tgfb1* gene deletion model.

**Figure 6.**
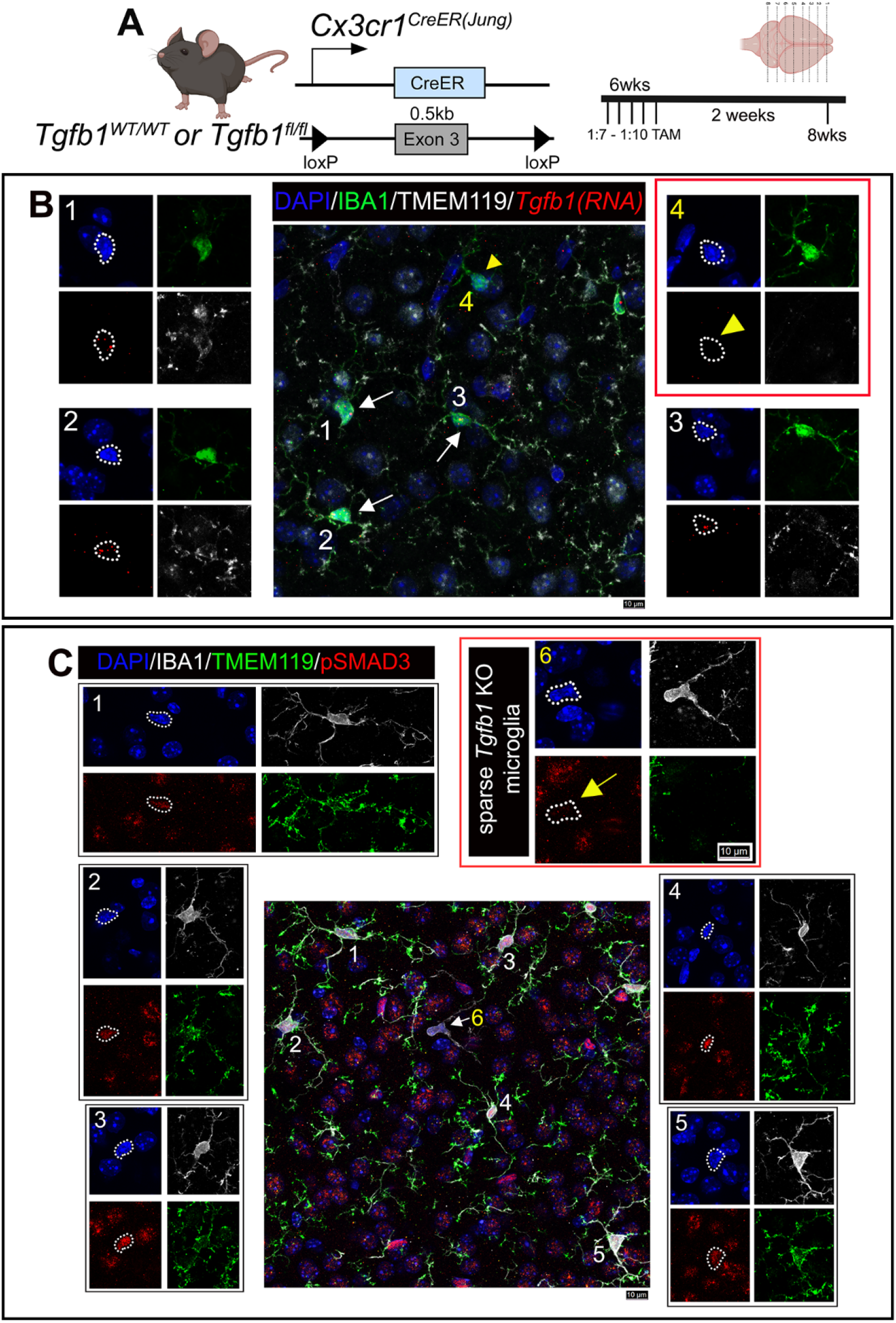
In-situ RNA-scope and IHC double labeling confirm loss of *Tgfb1* RNA and downregulation of TGF-β downstream signaling (pSMAD3) in dyshomeostatic individual microglia in the sparse *Tgfb1* iKO model. (A) The mouse model used to examine sparse iKO in microglia and experimental timeline with TAM dosage. (B) Representative image showing combined immunohistochemistry staining (for IBA1, TMEM119, DAPI) and *Tgfb1* RNA-scope hybridization. (B1-3) Surrounding normal microglia showing TMEM119 expression and *Tgfb1* RNA presence. (B4) A single microglia cell with loss of TMEM119 expression and loss of *Tgfb1* mRNA. White arrows were used to mark normal cells in the central panel. Yellow arrowhead used to mark individual iKO microglia. Note that tissue treatment for RNAscope analysis makes the IHC condition less ideal for morphology evaluation than regular IHC staining, however, IBA1 and TMEM119 expression are still distinguishable for individual WT or iKO microglia. (C) Representative image showing co-immunohistochemical staining with DAPI, IBA1, TMEM119, and pSMAD3. (C1-5) Surrounding normal microglia showing TMEM119 expression and pSMAD3 immunostaining. (C6) A single microglia cell with loss of TMEM119 expression and loss of pSMAD3 labeling. The yellow arrow (microglia #6) marks the individual iKO microglia. Representative results from n=3 iKO mice. Scale bar = 10µm. For additional representative images see Supplementary Figs 10 and 11.

### Loss of microglia-derived TGF-β1 ligand leads to transcriptomic changes indicating an activated state in both microglia and astrocytes in Cx3cr1-Tgfb1 iKO mice

To further characterize the transcriptional changes following the loss of microglial TGF-β1 ligand, microglia and astrocytes were sorted from the *Cx3cr1*^CreER(Jung)^*Tgfb1*^fl/fl^*R26-YFP* and *Cx3cr1*^CreER^ ^(Jung)^*Tgfb1*^wt/wt^*R26-YFP* animals 3 weeks after TAM administration based on YFP expression or ASCA2 immunolabeling (astrocyte staining, Supplementary Fig 12 and Fig 7)^38^ and subjected to RNAseq analysis. Purity of samples collected using this sorting method is validated by qRT-PCR for microglia and astrocytic signature genes respectively (Supplementary Fig 12D). We sorted brain microglia based on recombined YFP reporter expression (which labels about 90% parenchyma microglia in whole brain) instead of CD11b^+^/CD45^low^ to avoid the potential caveat that loss of TGF-β signaling in microglia may alter CD45 expression and selectively enrich for a subpopulation of microglia in the KO brain. PCA analysis shows that wildtype microglia samples and *Tgfb1* iKO microglia distinctively clustered together (Fig 7C). The heatmap shows significantly differentially expressed genes (fold change >= |1.5| and adj.pvalue < 0.05). In contrast to a recent study using Cx3cr1^CreER^*Tgfbr2*^fl/fl^ receptor inducible KO mice, which reported no changes in many homeostatic microglia signature genes in KO mice^9^, we observed a large set of differentially expressed genes (Heatmap Fig 7D) including downregulation of many microglia homeostatic signature genes (*P2ry12, Tmem119, Sall1*, etc. Fig7E, K) and upregulation of immune response regulating genes (*Tnf, Il1b*, Interferon responsive genes, Fig 7E, K). Using gene set enrichment analysis (GSEA) we observed upregulation of several pathways related to immune response, immune cell recruitment, and interferon response (Supplemental Fig 10B). We also observed downregulation in platelet aggregation pathway genes (Supplemental Fig 10C). For astrocytes, we observed upregulation of multiple A1-like genes (*Serping1, Ifit3, Gbp3*, Fig 7J, L) and interestingly we also observed an increased interferon response (*Irf7, Irf9*, Fig 7J). Consistently, GSEA analysis also showed increased interferon activity and decreased metabolic functions (NADH, Mitochondria, Acetyl Coa, Supplemental Fig 10D, E) which suggests a transition from metabolic support functions to an activated pro-inflammatory state in astrocytes from the MG-*Tgfb1* iKO brain. These data suggest that microglia and astrocytes had disrupted homeostatic functional activity after the loss of microglial TGF-β1.

**Figure 7.**
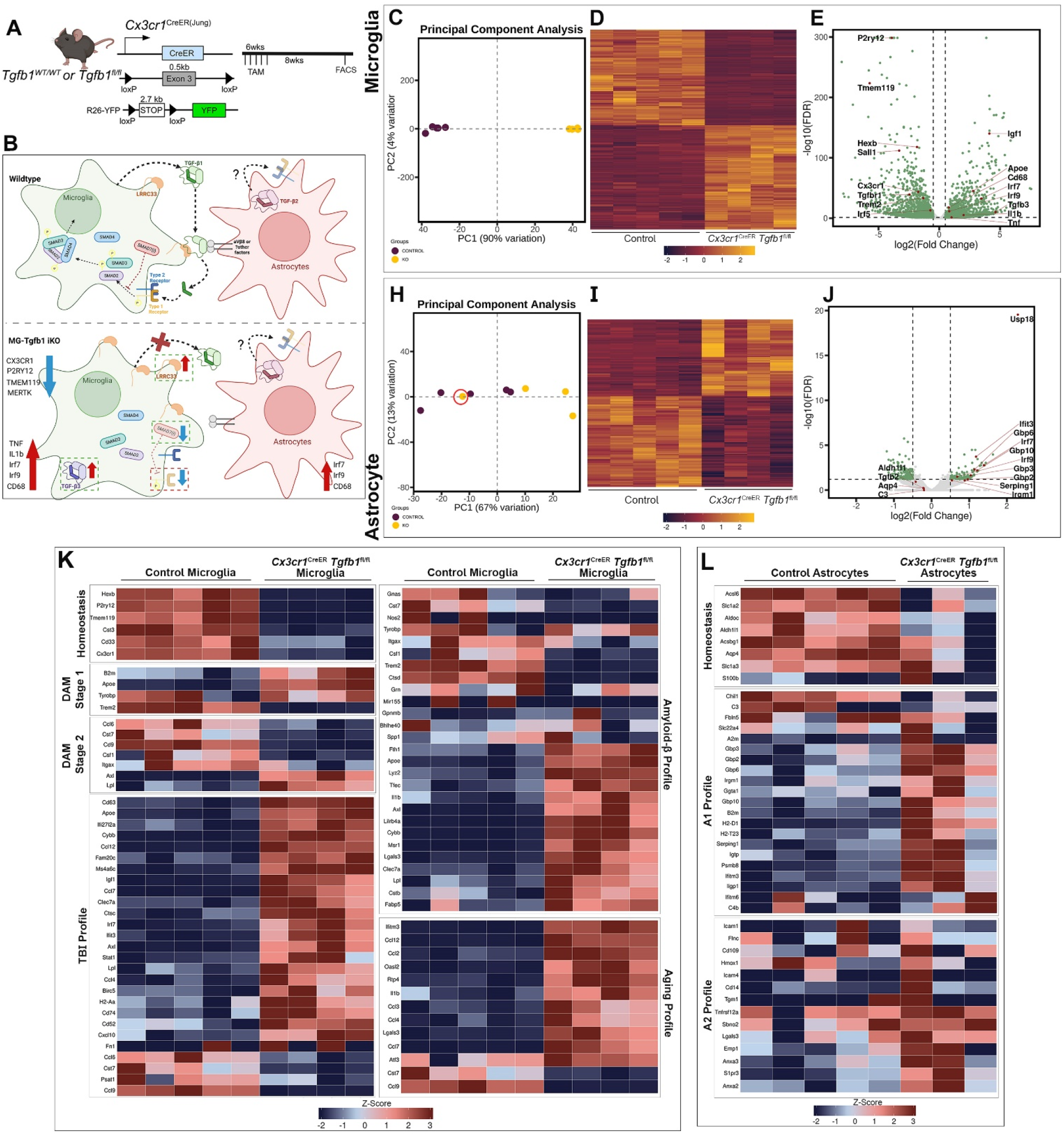
Transcriptomic analysis of microglia and astrocyte cells sorted from Cx3Cr1^CreER(Jung)^*Tgfb1*^fl/fl^ mice. (A) Mouse model used to induce *Tgfb1* KO and YFP reporter in microglia. (B) Summary of transcriptomic changes in microglia or astrocytes pertaining to both inflammatory responses and critical TGF-β signaling component genes. (C, H) PCA analysis plot of microglia and astrocyte samples. Note that one astrocyte sample from iKO mice clustered irregularly in the PCA plot which has an RNA Integrity Number (RIN) below 8 (red circle). (D, I) Heatmap of expression of significantly differentially expressed genes in microglia and astrocytes from control and iKO samples. (E, J) Volcano plot showing expression log fold changes. (K) Microglial differential gene expression observed across various gene sets including, homeostatic microglia genes^39–42^, stage 1 and 2 disease-associated microglia (DAM) genes^40^, injury exposed microglial (TBI)^42^, amyloid beta exposed microglia^39,41,42^, and aged microglia^41,42^. (L) Astrocytic differential gene expression was observed across different gene sets including, homeostatic astrocyte genes, A1, and A2 genes. Z-scores were calculated and plotted to display differential gene expression^43^. The astrocyte sample that had an RIN< 8 was excluded from this analysis.

We next wanted to compare the transcriptomic profile of the *Cx3cr1*^CreER^*Tgfb1*^fl/fl^ microglia and astrocytes in relation to previously characterized non-homeostatic microglial states. For microglia, we censured multiple previous studies to generate a list of signature genes associated with aging, CNS injury (Traumatic brain injury-TBI), and amyloid-beta pathological conditions^39–43^. We observed that three weeks after loss of microglial TGF-β1 ligand, microglia showed down regulation of microglial homeostatic genes (*Hexb, P2ry12, Tmem119, Cst3, Cd33, Cx3cr1*, Fig 7K) suggesting dyshomeostasis. We also observed increased expression of aging microglia signature genes (ex. *Ifitm3, Ccl12, Il1b Ccl2, Lgals3,* Fig 7K) and injury-associated TBI signature genes such as *Irf7, Igf1, Cxcl10, Ccl12, Axl, Cd63,* and *Cybb*. Recently amyloid beta-induced microglial transcriptomic changes have been profiled into DAM 1 and DAM 2 stages, the transition of which depends on TREM2 signaling^39–42^. Interestingly, *Tgfb1* iKO microglia resembles upregulation of a subset of amyloid beta-associated microglia profile genes, while showing down regulation of other amyloid beta profile genes. Upon further examination, we noted that the upregulated genes in iKO microglia represent DAM 1 signature genes (*B2m, Apoe, Tyrobp,* but a decrease in *Trem2 levels*) while downregulated genes in iKO microglia represent DAM 2 signature genes (*Ccl6, Cst7, Cd9, Csf1, Itgax*) which correlates well with the downregulation of TREM2 in iKO microglia. Additionally, after the loss of microglial TGF-β1, consistent with the observed reactivity in astrocytes by upregulation of GFAP protein, we observed downregulation of some astrocytic homeostasis genes (*Aldh1l1, Acsl6, Aldoc*), upregulation of A1 associated genes (*Gbp3, Gbp2, Gbp6, Psmb8,* Fig 7), and no discernable changes in the A2 associated genes (Fig 7L). This suggests that astrocytes might be adapting a neurotoxic-like (A1-like) rather than a neuroprotective (A2-like) profile.

Transcriptomic data from microglia and astrocytes also reveal interesting expression patterns of critical components of the TGF-β signaling pathways. Consistent with qRT-PCR data from sorted microglia and astrocytes (Supplementary Fig. 12D) and our data showing no observable changes following astrocytic TGF-β1 KO (Fig 3, Supplemental Fig 5, 6, & 7), RNA-seq data shows *Tgfb1*, *Tgfbr1*, *Tgfbr2*, and *Lrrc33* (a protein that is critical for latent TGF-β1 ligand activation) are all significantly enriched in microglia compared to astrocytes (% mRNA levels in microglia vs astrocytes: *Tgfb1*=500%, *Tgfbr1*=9500%, *Tgfbr2*=2600%, *Lrrcc33*=500%). Instead, astrocytes express *Tgfb2*, which is absent in microglia, suggesting *Tgfb2* might have a potential role in astrocyte function. Interestingly, we observed multiple compensatory mechanisms in response to MG-*Tgfb1* deletion: (1) an up-regulation of *Tgfb3* gene in microglia and no change of *Tgfb2* levels in astrocytes, (2) the upregulation of *Lrrc33* in microglia, a gene that has been demonstrated to be critical in activating the latent TGF-β ligand^16^, (3) down regulation of *Smad7*, which is a negative regulator of the TGF-β signaling pathway. Moreover, we observed a down-regulation of *Smad3* and *Tgfbr1* but not *Tgfbr2* mRNA, suggesting a TGF-β signaling-dependent feedforward regulation of *Smad3* and *Tgfbr1* expression (Fig 7B, Supplementary Fig 15).

### Loss of Microglia-derived TGF-β1 ligand leads to cognitive deficits in the Cx3cr1^CreER(Jung)^Tgfb1 iKO mice without affecting general locomotor function and motor learning

We next investigated whether the aging- and DAM-associated microglia profile and the induction of reactive astrocytes in the MG-*Tgfb1* iKO could affect neurological function in young adult mice. Full dosage TAM was used in this experiment to achieve maximum changes in microglia and astrocytes in the adult brain. A behavioral battery was used to examine general locomotion, motor coordination/learning, and cognitive function involving learning and memory. We first assessed voluntary movement in control and MG-*Tgfb1* iKO mice at 5 weeks after TAM injection using an automated open field locomotion tracking system and monitored mice for 23 hours with free access to food and water (Omnitech electronics INC, Columbus, OH). We did not observe any change in general locomotion in the *Cx3cr1*^CreER^*Tgfb1*^fl/fl^ +TAM animals compared to *Cx3cr1*^CreER^*Tgfb1*^wt/wt^ +TAM controls during the exploratory phase (1 hour after naïve exposure to the chamber), or during the light or the dark cycle (Fig 8K-N). Next, we carried out an acceleration rotarod test to evaluate motor coordination and motor learning. We specifically used a three-trial acceleration paradigm that starts at 1rpm and increases to 35rpm over the course of 5 minutes to evaluate their starting motor coordination, and how their performance improves over each trial. The *Cx3cr1*^CreER^*Tgfb1*^fl/fl^ + TAM animals did not show a difference in performance compared to control *Cx3cr1*^CreER^*Tgfb1*^wt/wt^ + TAM group in the rotarod test, suggesting that motor coordination and motor learning are not affected in the MG-*Tgfb1* iKO mice. However, interestingly, when we evaluated the cognitive function (spatial learning/memory) in control and MG-*Tgfb1* iKO mice using a 2-day Barnes Maze learning paradigm, *Cx3cr1*^CreER^*Tgfb1*^fl/fl^ TAM group showed an increase in latency to reach the escape hole and higher error trial numbers to locate the hole compared to the control mice, suggesting impaired spatial learning in the *Cx3cr1*^CreER^*Tgfb1*^fl/fl^ iKO mice. Importantly, *Cx3cr1*^CreER^*Tgfb1*^fl/fl^ mice that received vehicle treatment do not show any difference compared to control mice in any of the above behavioral tests (Fig 7C-J), demonstrating that the behavioral deficits in cognitive function measured by Barnes Maze in the *Cx3cr1*^CreER^*Tgfb1*^fl/fl^ + TAM mice are specifically caused by TAM-induced deletion of the microglial-*Tgfb1* gene in these mice. These data support that microglia-derived TGF-β1 ligand is critical in maintaining microglia homeostasis, astrocyte quiescence, and normal cognitive function in the adult brain.

**Figure 8.**
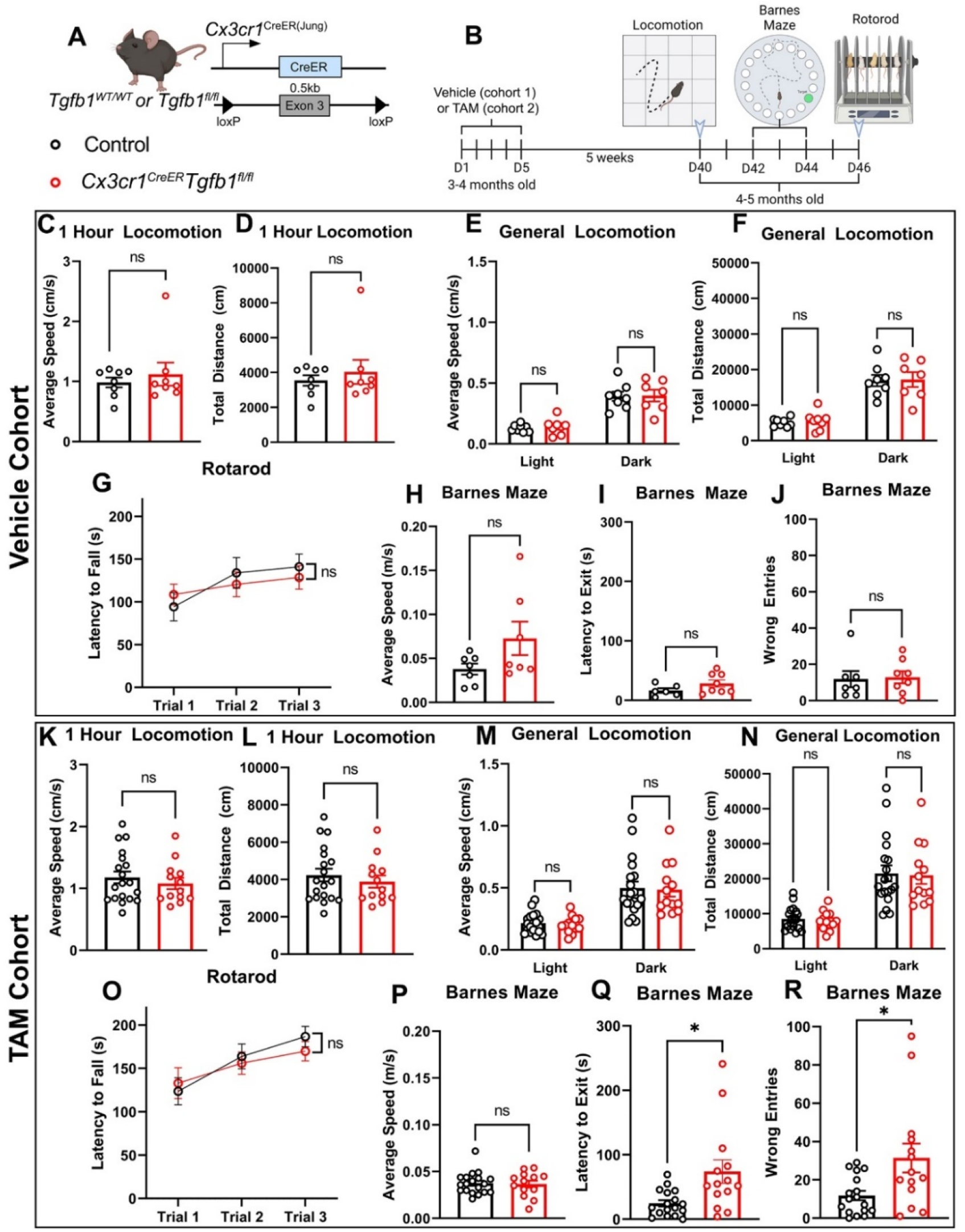
Behavioral assessment of full dosage Veh or TAM treated *Cx3cr1*^CreER^*Tgfb1*^wt/wt^ or *Cx3cr1*^CreER^*Tgfb1*^fl/fl^ mice to evaluate general motor function, motor coordination and learning, and learning and memory. (A) Mouse model used to induce *Tgfb1* KO in microglia. (B) Experimental timeline, showing the order of behavioral measurements. (C-J) Behavioral measurements in vehicle-treated *Cx3cr1*^CreER^*Tgfb1*^wt/wt^ or *Cx3cr1*^CreER^*Tgfb1*^fl/fl^ mice showing open field test (OFT) of the first hour in locomotion chamber (C) average speed and (D) total distance traveled. (E,F) average speed and total distance during the light and dark cycles in a 23-hour period. G) Accelerated rotarod learning test. (H-J) Barnes maze test showing (H) average speed during testing (I) latency to locating the target hole and (J) number of error trails before locating the target hole. (n=8 for each group). (K-R) Behavioral measurements from TAM-treated control and iKO mice showing open field test (OFT) of the first hour in locomotion chamber (K) average speed and (L) total distance traveled. (M, N) average speed and total distance during the light and dark cycles in a 23-hour period. (O) Accelerated rotarod learning test. (P-R) Barnes maze test showing (P) average speed during testing (Q) latency to locating the target hole and (R) number of error trails before locating the target hole. (control n=19, iKO n=13) (ns=not significant, * p<0.05, Student’s t-test)

## Discussion

Furthering our understanding of how TGF-β1 signaling is precisely regulated in the brain can provide important insight into microglia function during steady state and disease conditions. Our study addresses several critical gaps in our knowledge about CNS TGF-β ligand production and regulation and sheds light on how alteration of a single cytokine gene (*Tgfb1*) in microglia could causally contribute to cognitive deficits in young adult mice in the absence of brain injury or other disease model.

Currently, the prevailing understanding on the source of TGF-β1 ligand in the CNS has been speculated to be coming from multiple cell types and that TGF-β ligands can be widely shared among different cell types^11–14,30,31,37,44–47^. Several reviews have proposed the sharing of TGF-β1 amongst all the cell types, despite not yet having a well-rounded understanding of TGF-β1 ligand production and distribution^10–15^. CNS cell-type specific inducible TGF-β ligand gene deletion has not been studied previously and our study clarifies which cell types produce TGF-β1 ligands that are critical for maintaining microglia homeostasis in the adult mouse brain under physiological states. Our data support that microglia-produced TGF-β1 ligand is required for microglial homeostasis and subsequent astrocyte quiescence in the CNS. We used multiple microglia-CreER drivers to rigorously investigate this phenotype. Two independent *Cx3cr1*^CreER^*Tgfb1* iKO mouse lines both lead to a global loss of microglia homeostasis revealed by morphological changes and downregulation of homeostatic gene expression such as *Tmem119* and *P2ry12* without affecting serum or spleen levels of TGF-β ligand (demonstrated by both ELISA and FACS). We also show that astrocytic (via either inducible *Aldh1l1*^CreER^ line or constitutive postnatal deletion via the *mGfap*^Cre^ driver) or neuronal (via inducible *Camk2*^CreER^ line) deletion of the *Tgfb1* gene does not affect microglia morphology or expression of signature homeostatic microglia genes. Recent studies by us and others show that the *Cx3cr1*^CreER^ mouse lines^24,27^ recombine a portion of splenocyte macrophages even after the waiting period of >3 weeks, therefore, there is a possibility that the changes in microglia phenotype observed in the *Cx3cr1*^CreER^*Tgfb1* iKO mice could be due to parenchyma microglia population depletion and peripheral macrophage replacement in the brain. Alternatively, the activation of *Tgfb1* KO BAMs (which are also targeted by the *Cx3cr1*^CreER^ lines) could subsequently activate the rest of the parenchyma microglia. However, results from the *P2ry12*^CreER^*Tgfb1*^fl/fl^ mice and *Tmem119*^CreER^*Tgfb1*^fl/fl^R26-YFP mice also showed morphological and homeostatic microglial marker expression changes in the mosaic patches of YFP+ cells (indicating they were *P2ry12*+ or *Tmem119*+ at the time of TAM administration), supporting that the parenchymal resident microglia are altering their phenotype in response to deletion of *Tgfb1* gene, rather than being replaced or indirectly altered by peripheral monocytes or macrophages.

Additionally, by using low TAM dosage to achieve sparsely mosaic *Tgfb1* KO in very few individual microglia, our data supports that not only do microglia produce their own ligand to regulate their quiescent state, but they likely do so in an autocrine manner since *Tgfb1* gene deletion in sparsely distributed individual microglia leads to downregulation in TGF-β signaling (pSMAD3) and phenotypic changes in individual cells despite their surrounding WT microglia population. This data suggests that microglia regulate TGF-β signaling and related downstream pathways in a spatially precise manner which is consistent with the very low concentration of TGF-β1 ligand in brain tissue compared to spleen and serum levels. This mechanism is of particular importance in relation to disease or injury response since the glial activation cascade could be reliant on local fluctuating TGF-β1 levels. Importantly, TGF-β is synthesized as a latent form (L-TGF-β) whose activation requires release of the mature c-terminal domain from non-covalently bound latency-associated peptide (LAP)^16,48,49^. One recent study also suggested a possible highly localized mechanisms responsible for the releasing of the mature TGF-β1 ligand from the non-covalently bound prodomain (LAP) based on the coordinated molecular action of microglia-expressed LRRC33 (functioning as a LAP binding protein) and αVβ8 integrin^16^, possibly expressed on other cell types such as astrocytes.

While this microglial autocrine mechanism appears to be the primary mechanism for TGF-β1 signaling in microglia during homeostasis, interestingly, we observed that sparse individual KO microglia surrounded by WT microglia can recover to homeostatic state (measured by morphology and expression of TMEM119 and P2RY12) at 8 weeks after the loss of native TGF-β1 ligand production (Fig 5D). Note that on the populational level (with full TAM dosage and when most microglia are *Tgfb1* KO) at 8 weeks, microglia in the *Cx3cr1*^CreER^*Tgfb1* iKO mice still show morphological changes and loss of TMEM119 and P2RY12 expression. This suggests that the milieu environment of surrounding WT microglia is able to “reset” the sparse individual *Tgfb1* KO microglia in the sparsely mosaic KO mice but not when the majority of the microglia are KO cells that are activated. The mechanism for this recovery and to what extent the remaining “normal” microglia can help “reset” the mosaic activated microglia is not clear and warrants further investigation in future studies. Interestingly, our RNAseq data from the full dose TAM treated MG-*Tgfb1* iKO mice show that not *Tgfb2* but *Tgfb3* levels are upregulated in the MG-*Tgfb1* KO microglia (supplementary table 4), raising the interesting question of whether upregulated microglial TGF-β3 levels are able to compensate for the loss of TGF-β1 ligand in microglia. Previous studies have shown that *Tgfb1-* and *Tgfb3-*specific single KO mice have different phenotypes and that the swapping of the code sequence between Tgfb1 and Tgfb3 gene only leads to partial rescue of the phenotypes, suggesting non-overlapping functions of the two ligands, which is also supported by unique biophysical properties between the two ligands in recent studies^50^. Our results also suggest that upregulation of the TGF-β3 detected at 3 weeks post TAM in *Tgfb1*-KO microglia was not able to rescue the phenotype in microglia at up to 8 weeks post TAM in full dosage recombined mice. Interestingly, a recent study presented Cryo-EM structures which show that LRRC33 only presents L-TGF-β1 but not the -β2 or -β3 isoforms due to the differences of key residues on the growth factor domains^16^. This molecular selectivity offered by microglia-expressed LRRC33 could possibly explain why upregulated TGF-β3 expression in the TGF-β1 KO microglia could not compensate the loss of TGF-β1 ligand and rescue the phenotype in microglia in full TAM dosage MG-*Tgfb1* iKO mice. RNAseq data from sorted microglia and astrocytes from WT or MG-*Tgfb1* iKO brains also reveal interesting cell type specific transcriptomic regulation of the TGF-β signaling components in different cells during homeostasis or in response to the disturbance of TGF-β signaling. Specifically, we found that TGF-β1 is mainly enriched in microglia while TGF-β2 is enriched in astrocytes. Correspondingly, microglia express LRRC33 which preferentially presents TGF-β1 instead of TGF-β2 or -β3 for ligand activation. These patterns might explain why deletion of microglial TGF-β1 but not astrocytic TGF-β1 leads to the observed phenotypes in both microglia and astrocytes. Interestingly, loss of microglial TGF-β1 ligand also leads to down-regulation of *Smad3* and the TGF-βR1 but not TGF-βR2 in microglia, suggesting a feedforward regulation of TGF-β signaling on the expression of SMAD3 and TGF-βR1 which facilitates further TGF-β signaling. Conversely, *Tgfb1* KO microglia also upregulate *Tgfb3* and *Nrros* (LRRC33) while downregulating the inhibitory *Smad7*, reflecting an attempt to compensate for the loss of TGF-β signaling in KO microglia. These gene expression changes reflect a highly dynamic regulation of this critical signaling pathway and supports the precise spatially regulated autocrine mechanism in microglia. Consistently, none of these changes in the TGF-β signaling components are observed in astrocytes, indicating that the transcriptomic changes observed in astrocytes in the MG-*Tgfb1* iKO mice are likely not due to direct loss of TGF-β signaling in astrocytes. Whether genetic deletion of TGF-β2 or deletion of the type 1 or type 2 receptors in astrocytes would lead to activation of astrocytes warrants further investigation in future studies.

In the absence of the endogenous microglia-derived TGF-β1 ligand, microglia showed reduced ramification, decreased parenchymal homeostatic microglia signature gene expression, increased pro-inflammatory cytokine expression, and upregulation of interferon response genes. The expression profile of the *Tgfb1* iKO microglia aligned with damage associated microglia (DAMs)^40,42^, which have been described using both injury models (TBI)^42^ and disease states (amyloid beta pathology)^39–42^. Additionally, these transcriptomic profile changes also corresponded with observed gene expression changes in aging microglia^41,42^, suggesting that TGF-β1 signaling in microglia can provide vital insights into injury, neurodegenerative disease, and aging. The microglial *Tgfb1* iKO phenotype observed in our study is consistent with that described after constitutive *Tgfbr2* deletion in myeloid cells by the *Cx3cr1^Cre^* promoter during development^51^, which are more severe than the phenotype observed when *Tgfbr2* was deleted at P30^51^ or in a separate study at 2 months of age using the adult *Cx3cr1^CreER^Tgfbr2* inducible mice^26^. Factors such as the dosage and route of TAM treatment, efficiency of recombination of the floxed genes and whether cre-mediated recombination leads to complete absence of the target protein, or a truncated protein can all contribute to severity of the phenotypes. One specific potential caveat regarding the adult *Cx3cr1*^CreER^*Tgfbr2^fl/fl^*iKO study^9^ is that KO microglia were sorted from *Cx3cr1*^CreER(+/wt)^ mice which are heterozygous for the *Cx3cr1* gene while control microglia are sorted from *Cx3cr1*^CreER(wt/wt)^ mice which has both alleles of the *Cx3cr1* gene. Heterozygosity of *Cx3cr1* has previously been reported to cause changes in gene expression or function in microglia^52–56^ and therefore might introduce additional confounds to the data interpretation. Additionally, both previous studies^9,51^ used the same *Tgfbr2* floxed mouse line^57^ which has shown deletion of exons 2/3 do not alter the reading frame of the remaining exons leading to a truncated protein with normal serine/threonine kinase activity. This could potentially introduce an additional confound. In our study, we utilized a *Tgfb1* floxed mouse model (065809-JAX with 0.5kb of floxed region) which leads to a frame shift and results in complete absence of the active TGF-β1 ligand. This might explain the much more robust phenotypes in our adult iKO mice.

We also observed transcriptomic changes in astrocytes in the MG-*Tgfb1* ligand mice, featuring GFAP protein upregulation and increased interferon response genes. While GFAP is a pan-reactive marker, RNAseq data also shows upregulation of multiple A1-like neurotoxic astrocyte markers. Interestingly, deletion of *Tgfb1* gene in astrocytes via the constitutive *mGfap*^Cre^ driver line or the inducible *Aldh1l1*^CreER^ line did not induce morphological changes in microglia, nor GFAP expression in astrocytes, suggesting the activation of astrocytes in the MG-*Tgfb1* iKO mice is likely secondary to microglial profile change instead of direct loss of TGF-β signaling in astrocytes. This is consistent with the absence of *Tgfb1* gene in astrocytes which instead express *Tgfb2*. Loss of TGF-β signaling in microglia leads to upregulation of multiple pro-inflammatory cytokines which could in turn mediate the crosstalk between KO microglia and neighboring astrocytes. One such potential crosstalk can be through TNF signaling since it has been shown that TNF can promote A1-like astrocyte activation and the TGF-β1 ligand knockout microglia shows an increase in TNFα in KO microglia^58^.

Lastly, the behavioral analysis in the *Cx3cr1*^CreER(Jung)^*Tgfb1* mice shows that at 5 weeks following TAM treatment, there are significant deficits in the spatial learning and memory of MG-*Tgfb1* iKO mice without affecting the general locomotion function or motor learning. During the preparation of this paper, a recent study also reported a deficit in learning using a Morris Water Maze test in *Crybb1*^Cre^*Smad4* cKO mice^59^. The *Crybb1*^Cre^ driver targets embryonic macrophages (with some off-target recombination in OPCs and neurons) and therefore the observed learning deficits could result from a deficit of neurons and projections during development^59^. However, our data show that adult microglia critically rely on self-derived TGF-β1 ligand to maintain homeostasis and TAM-induced deletion of the TGF-β1 ligand in adult microglia leads to learning deficits in adult mice, suggesting an ongoing reliance on microglia-derived TGF-β1 ligand and TGF-β1 signaling in microglia to maintain normal cognitive function in adulthood. This result may have important implications for the role of microglial-TGF-β1 signaling in cognitive deficits observed during aging, neurodegenerative diseases, or after CNS injury. While constant basal TGF-β1 signaling is necessary for microglial homeostasis, TGF-β1 levels change with aging, injury, and disease^10,11,13,31^. Our results support that dysregulated microglia due to loss of TGF-β1 ligand with the transcriptomic features of DAMs and aged microglia might play a causal role in driving the cognitive deficits observed in these disease conditions and targeting TGF-β signaling might be a potentially therapeutic strategy to mitigate these deficits.

## Acknowledgement

Y.L is supported by NIH grants (R01NS125074, R01NS107365, R21NS127177). A.B is supported by NIH 1F31NS125930-01. I.I. is supported by NIH R35GM146890. We thank Chet Closson and the University of Cincinnati live imaging core (supported by NIHS10OD030402) for technical support. We also thank Dr. Xiang Zhang and the Genomics, Epigenomics and Sequencing Core at University of Cincinnati for RNAseq analysis support.

## Author contributions

YL conceptualized the study. YL, AB and EW designed the experiments. LM maintained all the mouse colonies and genotyped all mice in this study. AB performed TAM injection and the immunohistochemistry staining with help from MW and EW. AB performed all the behavioral analysis. MW carried out some of the microglia morphological analysis and astrocyte GFAP quantification. EW carried out the ELISA, microglia ablation experiment and qRT-PCR analysis of *Tgfb1* gene. AB performed cell sorting of microglia and astrocyte with assistance from the Flow Cytometry core at CCHMC and prepared all RNA samples for RNAseq analysis. The Genomics, Epigenomics and Sequencing Core at UC carried out the RNAseq. AP assisted the bioinformatics analysis of the RNAseq data in this study. AA and II carried out the flowcytometry analysis of *Tgfb1* expression in brain microglia and spleen myeloid cells. AB and YL drafted and revised the paper. All authors read, edited, and approved the final version of the manuscript.

## Supplementary material is available online

Table S1. Differentially expressed genes from sorted microglia collected using YFP reporter via FACS from *Cx3cr1*^CreER(Jung)^*Tgfb1^wt^*^/wt^ and *Cx3cr1*^CreER(Jung)^*Tgfb1*^fl/fl^. Related to Fig 7.

Table S2. Differentially expressed genes from sorted astrocytes collected using ASCA-2 antibody labeling via FACS from *Cx3cr1*^CreER(Jung)^*Tgfb1*^wt/wt^ and *Cx3cr1*^CreER(Jung)^*Tgfb1*^fl/fl^. Related to Fig 7.

Table S3. GSEA showing negatively regulated pathways from microglia from the RNA-seq data collected using *Cx3cr1*^CreER(Jung)^*Tgfb1*^wt/wt^ and *Cx3cr1*^CreER(Jung)^*Tgfb1*^fl/fl^. Related to Fig 7 and supplemental fig 13 and 14.

Table S5. GSEA showing negatively regulated pathways from astrocytes from the RNA-seq data collected using *Cx3cr1*^CreER(Jung)^*Tgfb1*^wt/wt^ and *Cx3cr1*^CreER(Jung)^*Tgfb1*^fl/fl^. Related to Fig 7 and supplemental fig 13 and 14.

Table S6. GSEA showing positively regulated pathways from astrocytes from the RNA-seq data collected using *Cx3cr1*^CreER(Jung)^*Tgfb1*^wt/wt^ and *Cx3cr1*^CreER(Jung)^*Tgfb1*^fl/fl^. Related to Fig 7 and supplemental fig 13 and 14.

Video S1. 3D rendered confocal image stack of coimmunostaining of IBA1 (red) and TMEM119 (white) in sparse MG-Tgfb1 iKO brain showing a single morphologically altered and IBA1+/TMEM119-KO microglia surrounded by normal neighboring microglia (IBA1+/TMEM119+).

## STAR Methods

### RESOURCE AVAILABILITY

#### Lead contact

Further information and requests for resources and reagents should be directed to and will be fulfilled by the Lead Contact, Yu Luo.

#### Materials availability

This study did not generate new unique reagents.

#### Data and code availability

- RNA-seq data have been deposited at GEO and are publicly available as of the date of publication. Accession numbers are listed in the key resources table.
- Microscopy data and behavioral test data reported in this paper will be shared by the lead contact upon request.
- No original code was generated in this study.
- Any additional information required to reanalyze the data reported in this paper is available from the lead contact upon request.

### EXPERIMENTAL MODEL AND SUBJECT DETAILS

#### Animals

The University of Cincinnati (UC) Animal Care and Use Program (ACUP) encompasses Laboratory Animal Medical Services (LAMS, animal facilities) and the Institutional Animal Care and Use Committee (IACUC) office. All animal protocols were approved by the IACUC. Mice were housed in the animal facility of University of Cincinnati on a 14-h light/10-h dark diurnal cycle. Food and water were provided *ad libitum*. The Cre-loxP recombination system was utilized to achieve cell-type specific constitutive or inducible knockout of *Tgfb1* gene. *Cx3cr* ^CreER(Jung)^ (JAX: 020940^27^), *Cx3cr1^CreER^*^(Littman)^ (JAX: 021160^24^) *P2ry12*^CreER^ (JAX: 034727^37^), *Tmem119*^CreER^ (JAX: 031820^36^), *Mgfap*^Cre^ (JAX:024098^33^), *Aldh1l1*^CreER^ (JAX:029655^32^), and *Camk2*^CreER^ (JAX:012362^35^) transgenic mouse lines in which the expression of Cre recombinase are under the control of the *Cx3cr1* (myloid cell), *P2ry12* (Microglia), *Tmem119* (Microglia and peri-vesicular fibroblast), mouse *Gfap* (astrocytes and adult neural stem cells), *Aldh1l1* (astrocytes), and *Camk2* (forebrain excitatory neurons) promoters, respectively were purchased from the Jackson Laboratory. These animals were crossed with mice carrying the floxed *Tgfb1* mouse line in which exon 3 of the *Tgfb1* gene is flanked by two loxP sites (JAX: 065809). Among all the mouse cre driver lines used in this study, the *Mgfap*^Cre^ mouse line is the only non-inducible and has previously shown to induce loxP specific gene recombination at perinatal stages in mice which targets a large percentage of astrocytes (>90%) and small percentage of cortical neurons (<1.3%) and some oligodendrocytes (<6% of total reporter positive cells) ^34^. All the other Cre driver lines have the CreERT2^60^.

#### Tamoxifen administration

Tamoxifen (TAM) injections were administered based on mouse body weight (BW). 180 mg/kg of BW was administered via oral gavage for 5 consecutive days for full dose induction. Mice were treated at 6-8 weeks of age with TAM. TAM cocktails were formulated from 100 μl EtOH and 900 μl of sunflower seed oil diluted for 30 mg of TAM powder. For sparse recombination, mice received 3 days of TAM at the dosage of 1:10 (18 mg/kg BW, we noticed that 1:7 dilution also give sparse labeled individual cells and similar results to 1:10. Each lab should test the titration of dilution in their own lab). Sparse TGF-B1 KOs were generated using a 1:7 -1:10 dilution of TAM in the vehicle (EtOH and sunflower seed oil) to achieve the desired dosage. Note that mice that receive the vehicle, diluted dosage (1:7-1:10) or full dosage of TAM should be housed separately to prevent TAM cross contamination between different groups.

#### Microglia ablation via PLX5622 Administration

Mice were treated with either PLX5622 diet (AIN-76A Rodent Diet With 1,200 PPM PLX5622, formulated by Research Diets with PLX5622 provided by Plexxikon) or the control diet (AIN-76A Rodent Diet, Research Diets, NJ). Animals had ad libitum access to the diet and water for the entirety of the study. For measuring *Tgfb1* mRNA levels after microglia ablation, C57bl6/J wildtype mice were treated with control or PLX5622 diet for 7 days. Brain tissue was harvested and processed for qRT-PCR as described below.

#### qRT- PCR

RNA was isolated from cortex using the RNAqueous-Micro Total RNA isolation kit (AM1931, ThermoFisher Scientific). CDNA was then generated using superscript III reverse transcriptase (18080044, ThermoFisher Scientific) or iScript cDNA Synthesis Kit (1708890, BioRad). The cDNA was then used for qRT-PCR using probes for *Hmbs1* (hydroxymethylbilane synthase), *Hprt1* (hypoxanthine phosphoribosyltransferase 1), *Iba1, Tgfb1, Alk5, Tgfbr2, Sall1, Glast, Glt*1, and *Atp1b2*. CDNA levels were quantified using a Roche Light Cycler II 480. Quantification of qRT-PCR values were normalized using the housekeeping gene Hmbs1 CT value, which did not change between groups after manipulation to account for potential variability in cDNA preparations.

#### ELISA

For ELISA analysis, tissue was collected after perfusion with phosphate buffer solution and flash frozen in cold isopropyl alcohol. Mouse serum is collected by clotted blood without any anticoagulant for 30 min followed by centrifugation at 1500 g for 10 min at 4°C. Serum is collected from the supernatant and frozen at -80C. The tissue was sectioned with a cryostat to punch 2mm punches of tissue. Tissue was placed in RIPA buffer then homogenized using sonication at 30% amplitude, for 3 second pulses with 2 second pauses. BCA method was used to determine total protein concentration in the samples and Quantikine ELISA Human TGF-β1 kit (R&D Systems, Minneapolis, MN) was used to analyze TGF-β1 ligand levels following the instruction from the manufacturer.

#### Tissue collection for Flow Cytometry or FACS

Mice were transcardially perfused with cold 1x HBSS for 2-3 minutes. The brains and spleens were extracted and mechanically dissociated with a scalpel before using the papain dissociation kit (9001-73-4, Worthington Biochemical Corporation). For spleens, following dissociation, red blood cells were lysed using ammonium chloride. For the brains, once dissociated, cells were suspended in a 37% percoll solution and spun at 800g for 20 minutes to remove excess myelin and debris. The cells were collected, washed, resuspended in FACS buffer containing PBS with 1% (v/v) fetal bovine serum and 0.1% (w/v) NaN3 (Sigma), and counted. The number of cell subpopulations in the CNS were determined by multiplying the percentage of lineage marker–positive cells by the total number of mononuclear cells isolated from the brain. Transcriptional and translational inhibitors actinomycin, anisomycin, and typtolide were used to prevent activation of microglia during the preparation of tissues as was previously described by Marsh et al.^61^. Inhibitors were added to the dissection solution and the papain enzyme cocktail from the Worthington kit.

#### Flow Cytometry Analysis of TGF-β1 Expression

To carry out flow cytometry analysis, the Fc receptors were initially blocked using anti-mouse CD16/32 (0.25 μg; ThermoFisher) for 15 min at 4°C. Cells were then washed with FACS buffer and stained for surface marker for 30 min at 4°C using the specified antibodies. These antibodies included: CD45 (clone 30-F11), CD11b (clone M1/70) and TGF-β1 (clone TW7-16B4) (all from Biolegend). Cells were then washed with PBS and viability staining was performed using the LIVE/DEAD fixable dead cell stain kit (Invitrogen). Following viability staining, cells were washed with PBS and resuspended in FACS buffer for Flow cytometry analysis. Cells were acquired on a BD Canto II and analyzed using FlowJo X software (vX10). As controls, fluorescence minus one (FMOs) were used to place the gates for analysis. For flow cytometry analysis, cells were first gated according to FSC-SSC, then restricted to singles cells and live cells. Myeloid cells were identified as CD45+ CD11b+.

#### FACS of microglia and astrocytes for qRT-PCR and RNA-seq

Gating was determined using the yellow fluorescent protein expressed by *Cx3cr1*^CreER^-R26-YFP for microglia collection, and ASCA-2 APC conjugated antibody (130-117-535, Miltenyi Biotec) for astrocytes. Any double positive cells were excluded from the gating to improve the purity of samples.

#### Bulk RNA-Sequencing

Non-directional RNA-seq was performed by the Genomics, Epigenomics and Sequencing Core at the University of Cincinnati. To summarize, the quality of total RNA was QC analyzed by Bioanalyzer (Agilent, Santa Clara, CA). About 100 pg total RNA was used as input for cDNA amplification using NEBNext Single Cell/Low Input RNA Library Prep Kit (NEB) under PCR cycle number of 15. After Bioanalyzer QC, 20 ng cDNA was used for library construction under PCR cycle number of 6. After library QC and quantification via Qubit quantification (ThermoFisher, Waltham, MA), individually indexed libraries were proportionally pooled and sequenced using NextSeq 2000 Sequencer (Illumina, San Diego, CA) under the sequencing setting of PE 2×61 bp to generate about 60M reads. Once the sequencing was completed, fastq files were generated via Illumina BaseSpace Sequence Hub.

#### RNA-sequencing Analysis

RNA-seq reads with adapter sequences or bad quality segments were trimmed using Trim Galore! v0.4.2^62^ and cutadapt v1.9.1^63^.The trimmed reads were aligned to the reference mouse genome version mm10 with STAR v2.6.1e^64^. Duplicated aligned reads were removed using sambamba v0.6.8^65^.Gene-level expression was assessed by counting features for each gene, as defined in the NCBI’s RefSeq database^66^. Read counting was done using featureCounts v1.6.2 from the Rsubread package^67^. Raw counts were normalized as transcripts per million (TPM).Differential gene expressions between groups of samples were assessed with R package DESeq2 v1.26.0^68^. Gene list and log2 fold changes are used for GSEA^69,70^ analysis using GO pathway dataset.Plots were generated using the ggplot2^71^ package and base graphics in R. The PCA analysis comparing astrocytes from control samples and astrocytes from MG-*Tgfb1* iKO mice show one astrocyte sample from iKO mice diverge from other iKO samples and this sample had a lower RNA integrity number (below 8 while all other samples have RIN of >8), suggesting partial RNA degradation. We included this sample in the PCA plot, general DEG heatmap, and volcano plot, however this sample was excluded for characterizing astrocytic activation profile.

#### Immunohistochemistry

Mice were perfused with 4% PFA and drop fixed overnight before being transferred to 20% sucrose, followed by 30% sucrose once the tissue had sunk. Then tissue was sectioned on a cryostat in 30 μm thickness prior to implementing IHC. Antibodies for GFP (1:1000, Invitrogen or 1:500, Aves), IBA1 (1:500, Abcam), IBA1 (1:1000, Wako), P2RY12 (1:500, Biolegend), TMEM119 (1:2000, GeneTex), GFAP (1:1000, Sigma), NEUN (1:1000, Biolegend), CD68 (1:4000, BioRad), and pSMAD3 (1:150, Abcam) were used. Tissue was blocked for 1 hour at room temperature (RT) in 4%BSA/0.3% Triton-X100, then incubated overnight at 4C in primary antibody. Tissue was then incubated for 2-3 hours in appropriate secondary antibodies conjugated with Alexa fluorescence 488, 555, 647, or 790.

#### RNA-Scope

To fluorescently label RNA, RNA-scope was employed using the ACD RNA-scope Multiplex v.2 kit. Samples for RNA-seq were perfused with cold PB and the brain were dissected out and drop fixed in 4% PFA for 7 hours before transferring to 20% and 30% sucrose. Brains were sectioned on a Leica Cryostat at the thickness of 16um and directly mounted onto Superfrost Plus glass slides. RNAscope hybridization steps were carried out following the instruction from the manufacture. A *Tgfb1* probe was used (443571, ACD Biosciencse) to label ligand RNA and an ALK5 probe was used (406201, ACD Biosciences) to label the TGF-β1 type I receptor. Immunohistochemistry was carried out after the RNAscope hybridization as described above to identify different cell types or expressions of different homeostatic microglia markers.

Confocal images were obtained for IHC and RNAscope/IHC samples at the University of Cincinnati Imaging Core utilizing the Leica Stellaris 8 Confocal microscope. 3D reconstruction of the Z-stacks was performed in LAS X software. Imaging was also analyzed using Neurolucida image analysis program (MBF Bioscience) for detailed morphological analysis.

#### Image analysis

NIS-Elements was used for microglial morphology quantification. A trace function was utilized for measuring the microglia process length (μm) and a counting function was utilized to quantify the number of terminal ends for each microglial process. Reactive astrocytes were quantified by measuring the area of GFAP immunoreactive astrocytes in the field via thresholding against the background. CD68 quantification was accomplished through the image processing package of ImageJ, Fiji. Z-stacks from confocal imaging were merged into a max projection on the Leica Application Suite X (LASX), and then exported to Fiji. The threshold function was utilized on the fluorescent staining of CD68 against the background, and averages were taken of the mean area. On average 4-6 images from 3-4 brain sections at similar brain regions were analyzed per mouse and the value from multiple images were averaged for each mouse which is used as a single data point in statistical analysis. Neurolucida (MBF Bioscience) was utilized for 3D microglial reconstructions. Microglia cell bodies were constructed by tracing the outer perimeter with the cell body trace function each time a primary microglia process branched out. Then each process was traced with the process tracer function. For pSMAD3 quantification, ImageJ was used to identify IBA1+ and IBA1-nuclei, the nuclei were then traced, and the median fluorescent intensity was measured. Multiple microglia (7-10 microglia) were analyzed from randomly sampled multiple images in individual mice and the average from multiple cells is used as a single data point in statistical analysis.

#### Behavioral Assays

*Locomotion* To measure general locomotion function, automated 42 × 42 × 31 cm Plexiglas open box for 23 hours with ad libitum food and water (on the lid) and fresh bedding on the chamber of the floor (Omnitech electronics INC, Columbus, OH) with laser sensors were used to monitor animal behavior. Animals were housed in a dual chamber, allowing for scent exchange but not physical interaction. Lighting was set to mimic that of the animals normal housing rooms, on a 14/10hr light/dark cycle. Data was automatically scored using Fusion (Accuscan, Columbus, OH, USA).

#### Rotarod

Using the Roto-rod Series 8 apparatus (IITC Life Science Inc., Woodland Hills, CA) mice were given an acceleration paradigm. The rods rotation started at 1RPM and reached 30RPM by the end of the 5-minute session. The test concluded when the animal fell from the rod but were not returned to the home cage until all animals finished the test. Animals performed 3 trials and were allotted at least 15 minutes in their home cage before starting their next trial. 70% ethanol was used to clean the apparatus in between animals.

#### Barnes Maze

Barnes Maze apparatus (Stoelting company, Wood Dale, IL) was designed as a gray circular platform with 91 cm in diameter, 90 cm in height with 20 5cm diameter holes equally distributed around the edge of the platform. Of the 20 holes, 19 had 2cm deep gray trays beneath, with one of the holes having a 5cm deep escape box. A short challenging Barnes maze learning paradigm was used to assess spatial learning and memory. Animals were given 2 training sessions, 4 hours apart, with each training session ending when the animal located and fully entered the escape box. LED lights, a heat lamp and fan were used to motivate escape behavior. The following day (24 hours later) a test session occurred, with the test ending when the animal fully entered the escape box. In-between animals the maze was sanitized using 70% ethanol. Data was automatically scored using AnyMaze (Stoelting company, Wood Dale, IL).

#### STATISTICAL ANALYSIS

All studies were analyzed using SigmaPlot. Results are expressed by mean±SEM of the indicated number of experiments. Statistical analysis was performed using the Student’s t-test, and one- or two-way analysis of variance (ANOVA), as appropriate, with Tukey post hoc tests. A p-value equal to or less than 0.05 was considered significant.). Graphs were made in GraphPad Prism and some portions of figures were generated with Biorender.com.

## Key Resources Table

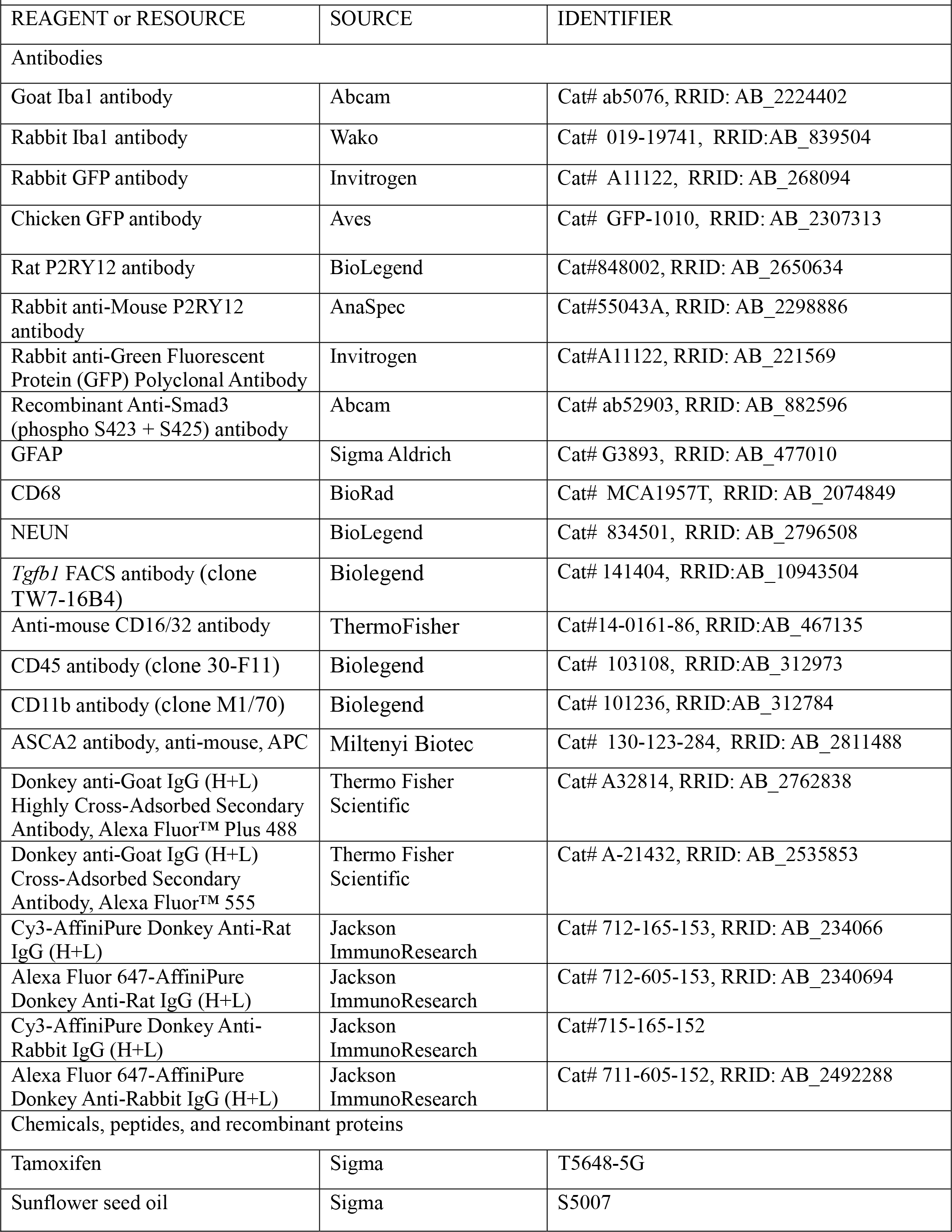
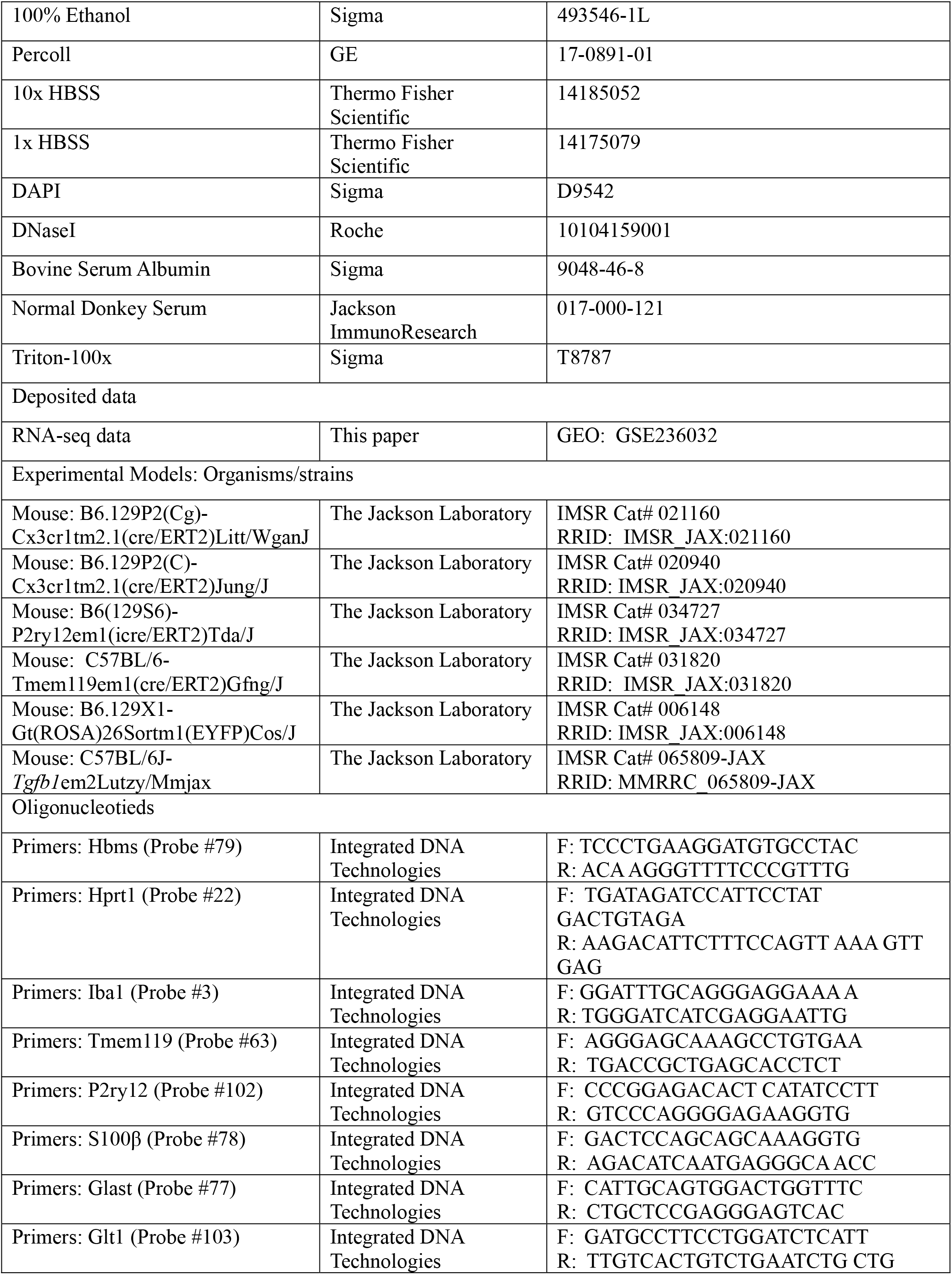
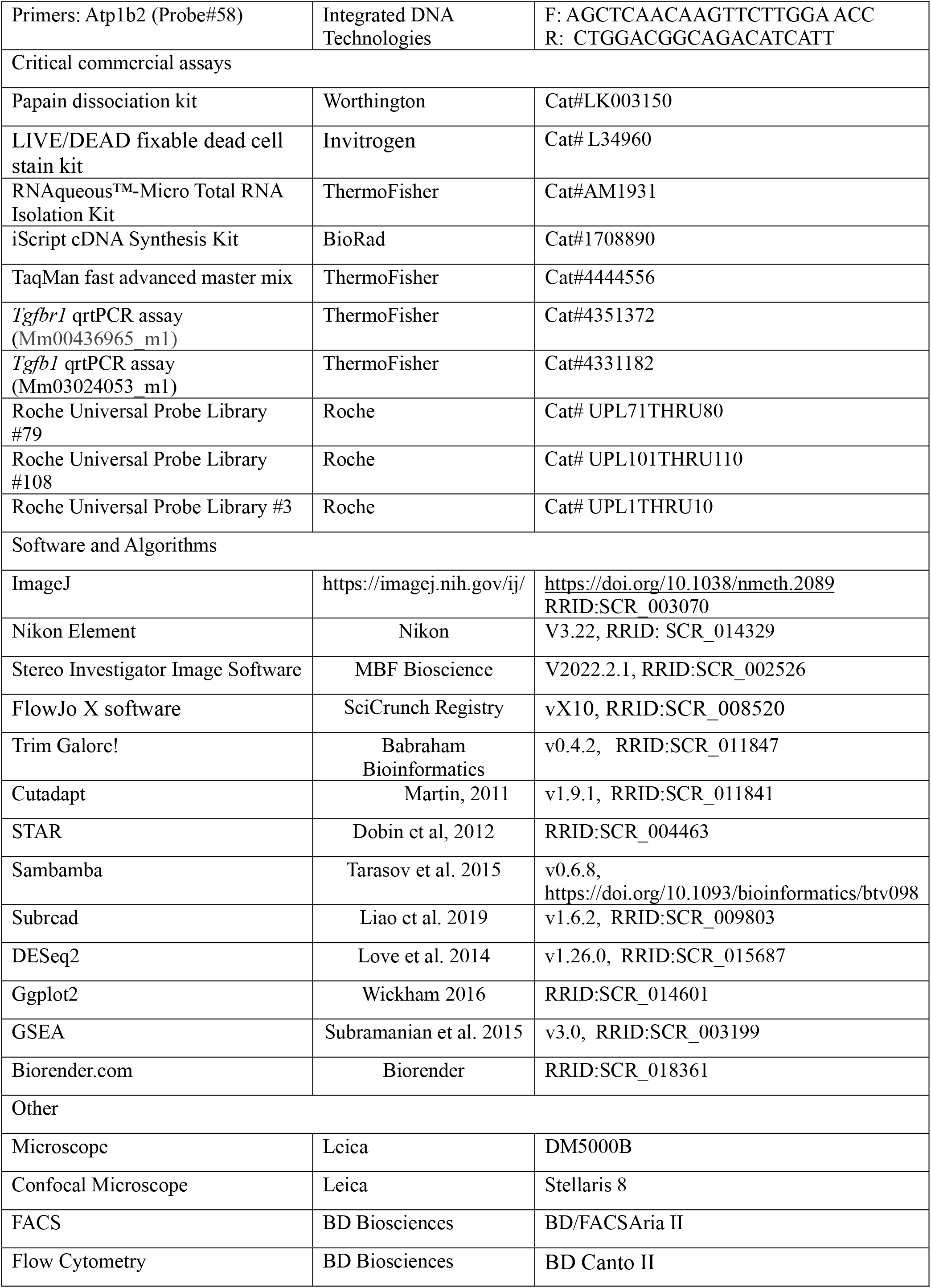

---

**Supplementary Figure 1.**
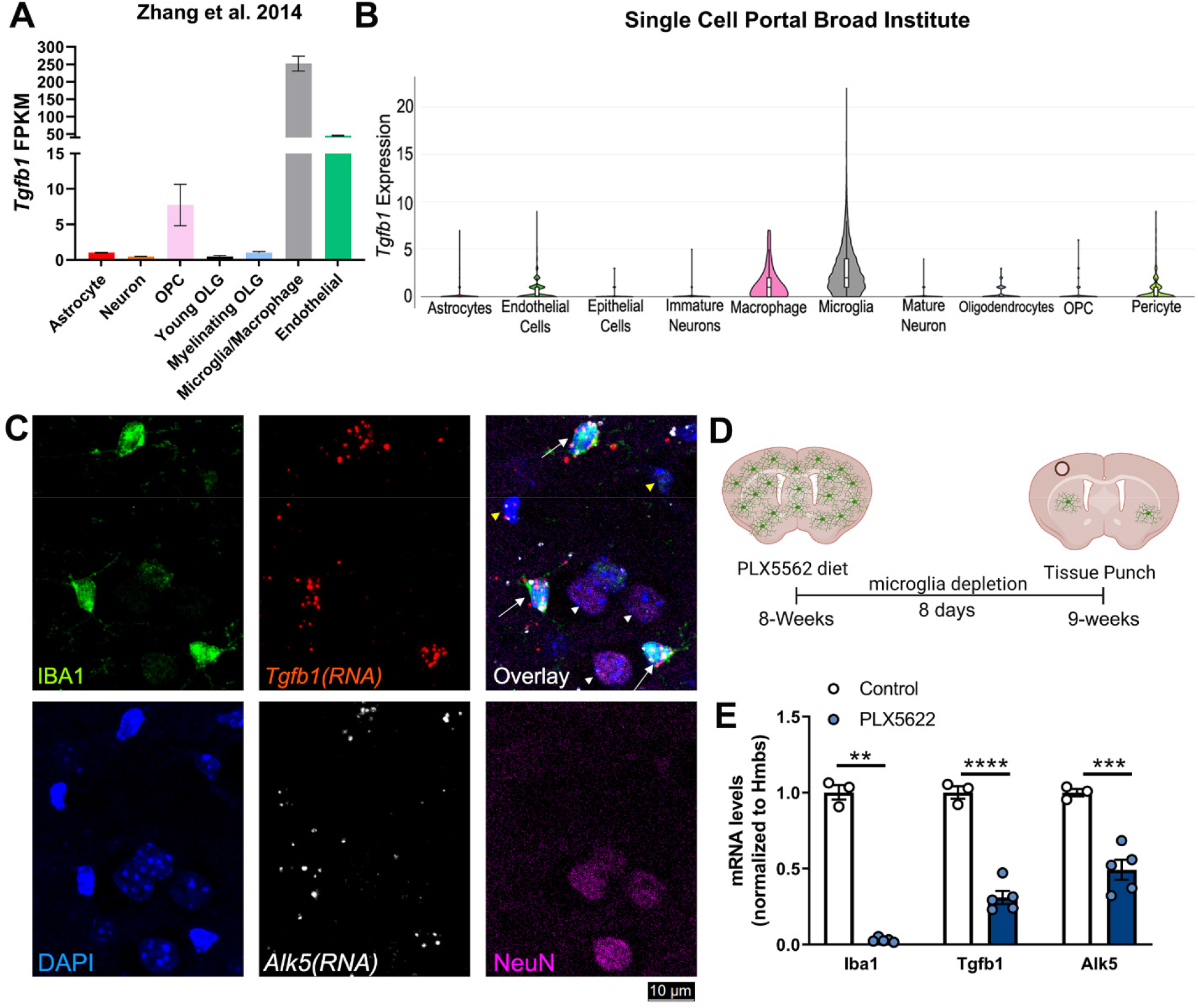
*Tgfb1* gene expression is enriched in microglia in adult mouse brains. (A, B) Publicly available single-cell RNA-sequencing showing microglial enrichment for *Tgfb1* expression from Zhang et. al 2014 (A) showing fragments per kilobase of transcript per million reads mapped (FPKM) and a second independent Single-cell portal data set^1^ (B) showing similar microglial enrichment of the *Tgfb1* expression among different cell types. (C) Representative images using combined RNA-scope/Immunohistochemistry visualizing RNA-scope in situ signal for *Tgfb1* mRNA and *Tgfbr1* (*Alk5)* mRNA combined with IHC for IBA1 and NEUN showing colocalization of *Tgfb1 and Alk5 mRNA with IBA1+ microglia (white arrow) but not in NEUN+ neurons (white arrowhead). Note some Tgfb1 mRNA signals in IBA1-cells (yellow arrowhead).* (D) Experimental timeline for PLX5562 ablation of microglia, (E) microglia ablation leads to a significant decrease of *Iba1*, *Tgfb1*, and *Alk5* mRNA levels (normalized to Hmbs). Mean±SE, ** = p<0.001, *** = p<0.0005, **** = P<0.0001, Student’s t-test (each data point represents an individual animal).

**Supplementary Figure 2.**
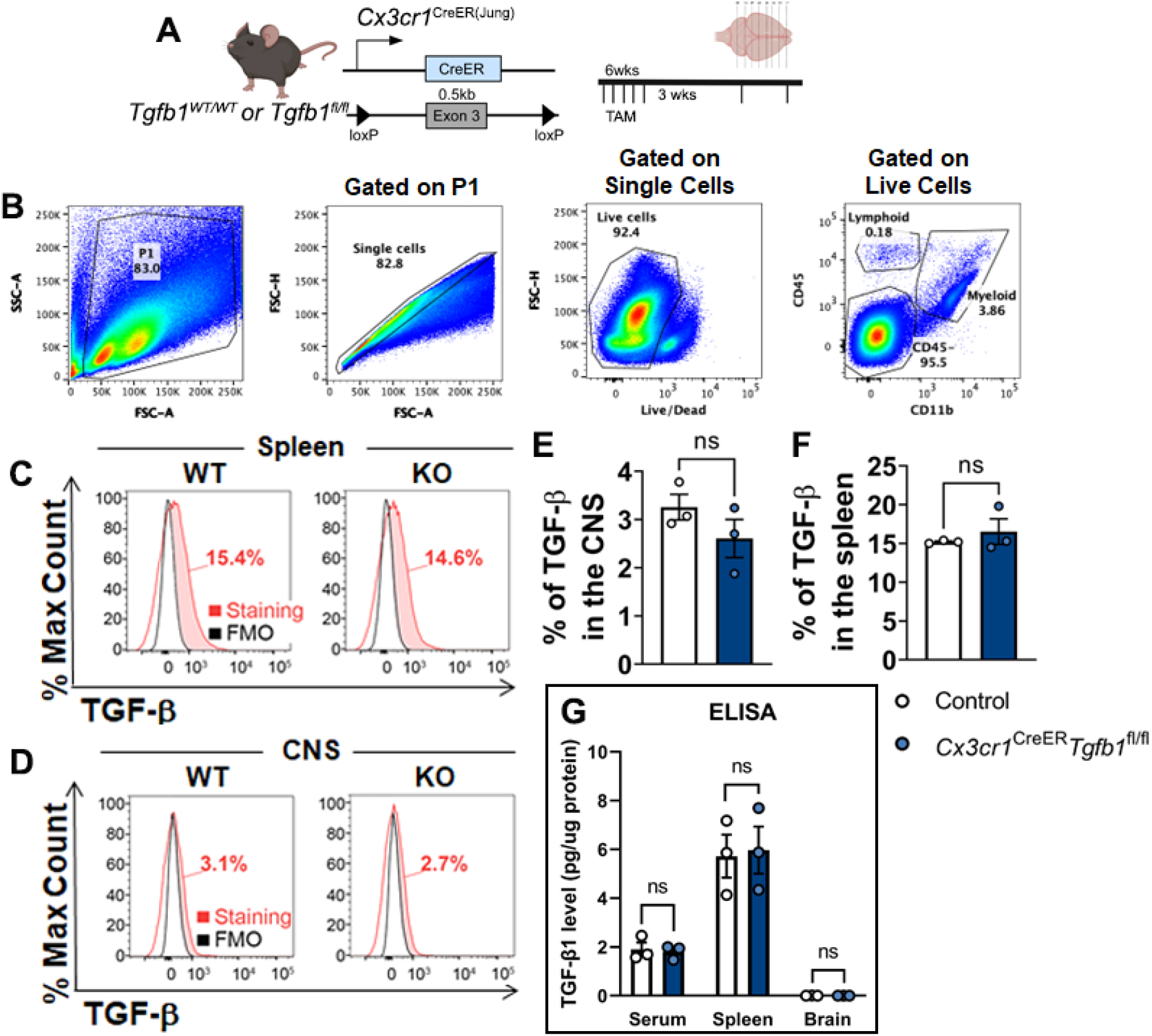
FACS or ELISA detection of TGF-β1 protein in the brain, serum, and spleen in WT or MG-*Tgfb1* iKO mice. (A) Timeline for using Cx3cr1^CreER^ driver to target myeloid cell inducible *Tgfb1* loss (samples were harvested at 3 weeks post TAM). (B) Gating strategy for identifying the myeloid cells in single-cell suspension. Cells were first gated according to FSC-SSC, then restricted to singles cells and live cells. Myeloid cells were identified as CD45+ CD11b+. (C, D) FACS analyses of TGF-β expression by myeloid cells in the spleen (top histograms) and in the CNS (lower histograms). The fluorescence minus one (FMO) is represented in black histograms and TGF-β immunostaining is shown in red. TGF-β staining above the background (FMO) is shown in solid red and represents the percentage shown for each analysis. (E, F) Compilation of TGF-β expression by flow cytometry on myeloid cells from the spleen and CNS of WT and KO mice. (G) ELISA quantification (pg/µg total protein) from serum and tissue from the spleen and brain (n = 3 mice per group) showing no difference in TGF-β protein levels in serum or spleen of Cx3cr1CreER-*tgfb1* iKO or control mice and that the brain TGF-β ligand levels are below the detection limit of the kit. Mean±SE, Student’s t-test (each data point represents a single animal).

**Supplemental Figure 3.**
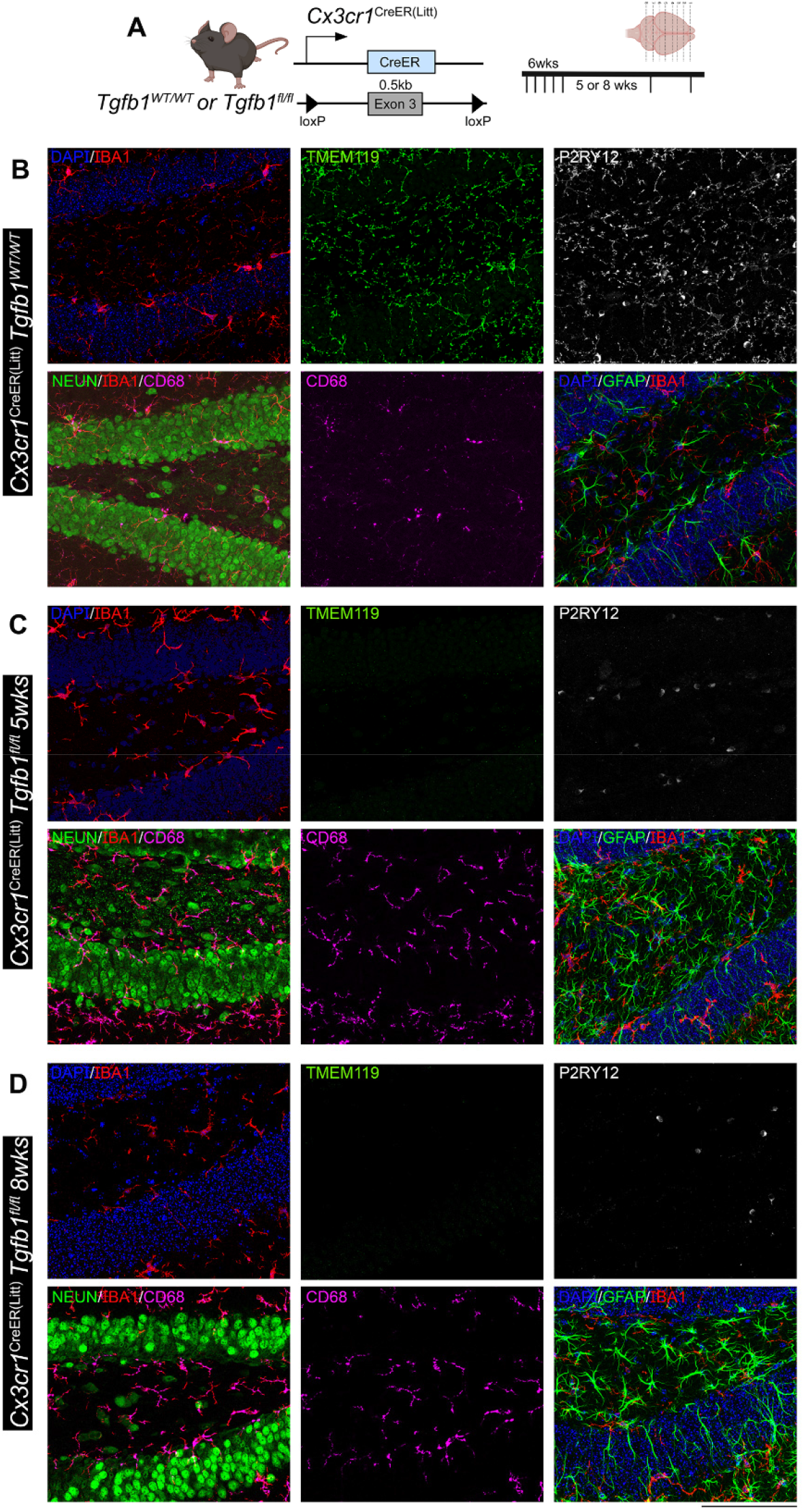
Microglia-specific *Tgfb1* gene deletion results in loss of homeostasis of microglia and increase in reactive astrocytes in the hippocampus of the adult mouse brain. (A) Mouse model for targeting microglial *Tgfb1* and timeline. (B) Representative immunohistochemistry images of IBA1, TMEM119, P2RY12, CD68, NeuN, and GFAP in the hippocampus of (C) Control animals, (D) *Cx3cr1*^CreER(Litt)^*Tgfb1*^fl/fl^ knockouts 5 weeks after tamoxifen administration, and (E) *Cx3cr1*^CreER^ ^(Litt)^*Tgfb1*^fl/fl^ knockouts 8 weeks after tamoxifen administration. Representative results from n=3-5 mice/group, Scale bar = 100µm.

**Supplemental Figure 4.**
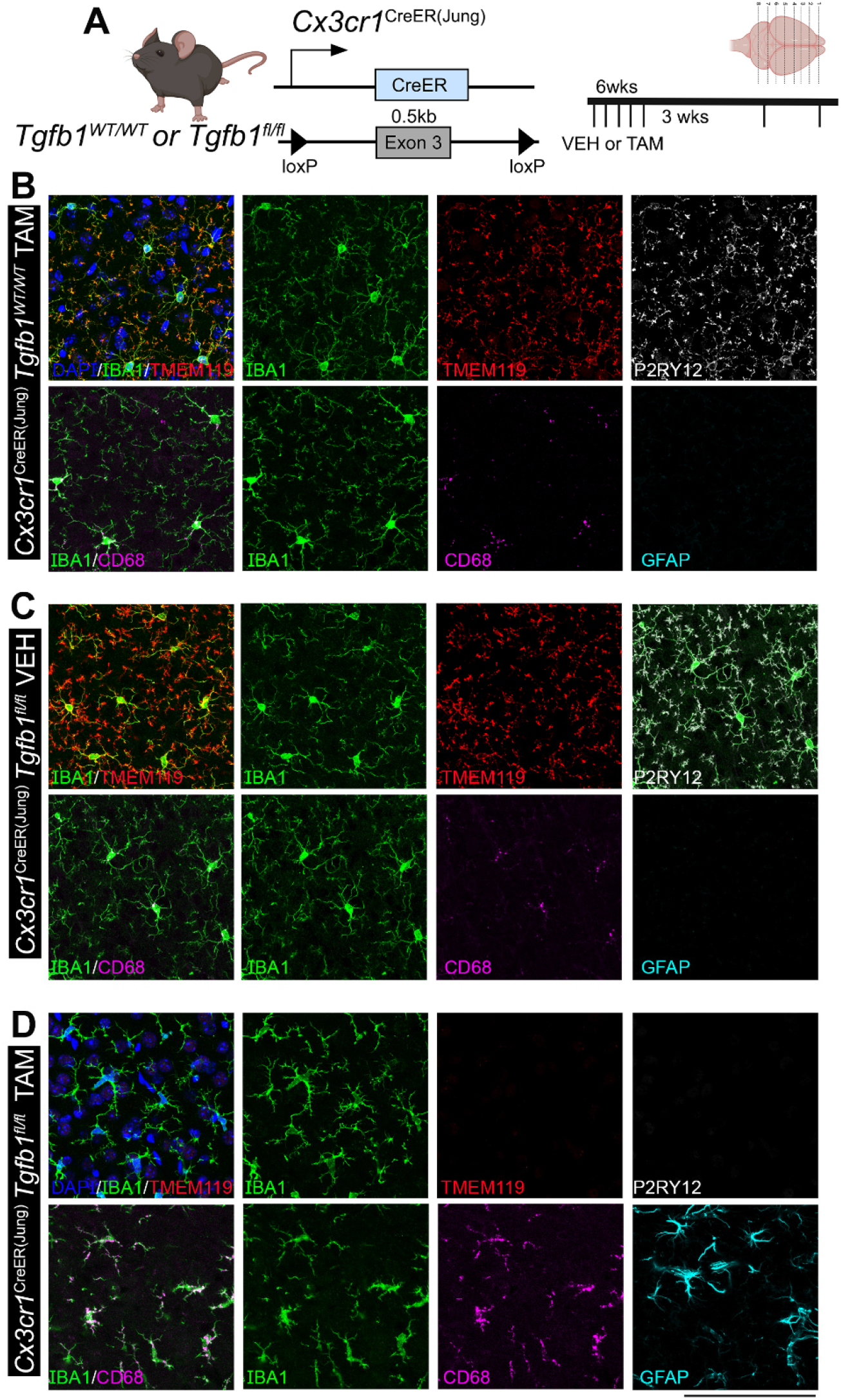
Microglia specific *Tgfb1* gene deletion in a second independent Cx3cr1^CreER(Jung)^ driver line results in a similar phenotype of loss of homeostasis of microglia and an increase in reactive astrocytes in the cortex of adult mouse brain. (A) Mouse model for targeting microglial *Tgfb1* and timeline. (B-D) Representative images for immunohistochemistry looking at IBA1, TMEM119, P2RY12, CD68, and GFAP in (B) Control Cx3cr1CreER(Jung) tgfb1 wt/wt +TAM animals, (C) *Cx3cr1*^CreER(Jung)^*Tgfb1*^fl/fl^ mice + Veh control and (D) *Cx3cr1*^CreER(Jung)^*Tgfb1*^fl/fl^ mice + TAM at 3 weeks after tamoxifen administration. Representative results from n=3-5 mice/group, Scale bar = 100µm.

**Supplemental Figure 5.**
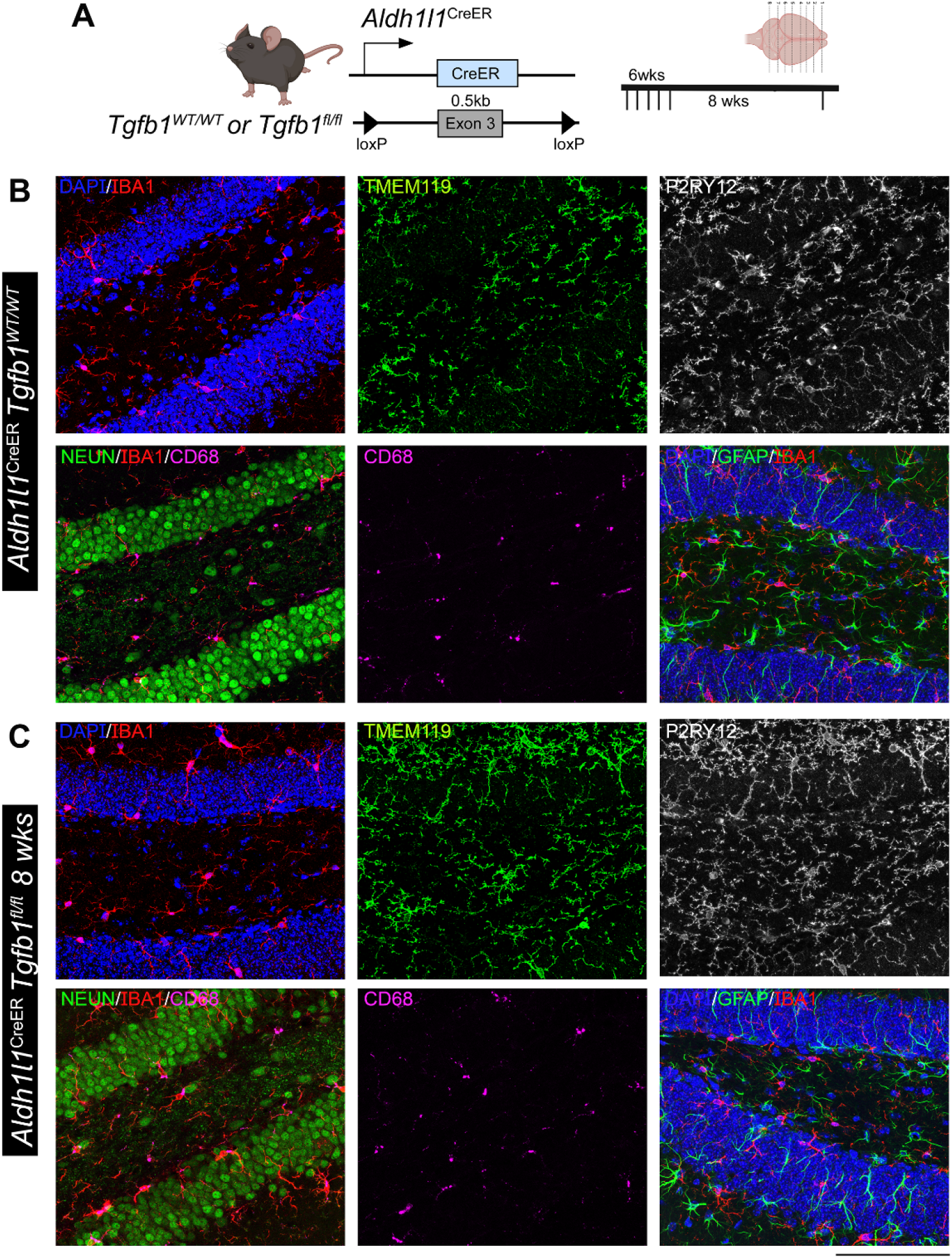
Astrocyte-specific *Tgfb1* gene deletion in the *Aldh1l1*^CreER^ driver does not affect the homeostasis of microglia or GFAP expression in astrocytes in adult mouse brain (hippocampus). (A) Astrocyte iKO mouse model and experimental timeline. (B,C) Representative immunohistochemistry images of hippocampus from TAM treated (8 weeks post) control (B) *Aldh1l1*^CreER^*Tgfb1*^wt/wt^ and (C) iKO *Aldh1l1*^CreER^ *Tgfb1*^fl/fl^ tissue showing IBA1, TMEM119, P2RY12, CD68, and GFAP immunostaining. Representative results from n=3-5 mice/group, Scale bar = 100µm.

**Supplemental Figure 6.**
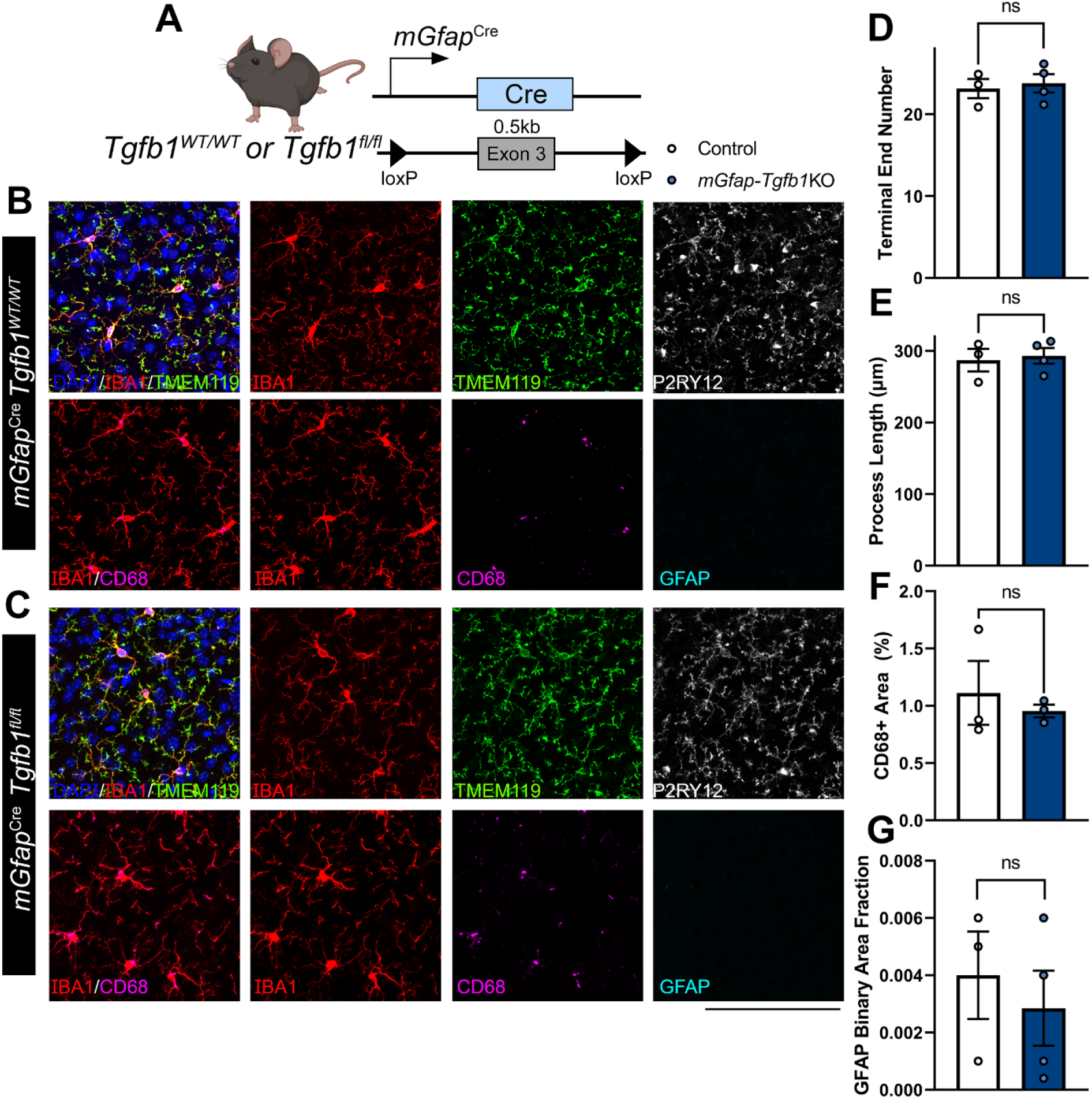
Astrocyte-specific *Tgfb1* gene deletion in the perinatal constitutive mGfap^Cre^ driver line does not affect the homeostasis of microglia or GFAP expression in astrocytes in adult mouse brain (cortex). (A) Astrocyte constitutive KO mouse model used and experimental timeline. (B,C) Representative images of control *mGfap*^Cre^*Tgfb1*^wt/wt^ (B) and cKO *mGfap*^CreER^ *Tgfb1*^fl/fl^ tissue showing IBA1, TMEM119, P2RY12, CD68, and GFAP immunostaining. Quantification of microglia ramification via (D) terminal end number, (E) process length, and (F) CD68+ immunoreactive % area. (G) Quantification of astrocyte reactivity using GFAP+ immunoreactive area fraction. (control n=3: cKO n=4) ns=not significant. Mean±SE, Scale bar = 100µm.

**Supplemental Figure 7.**
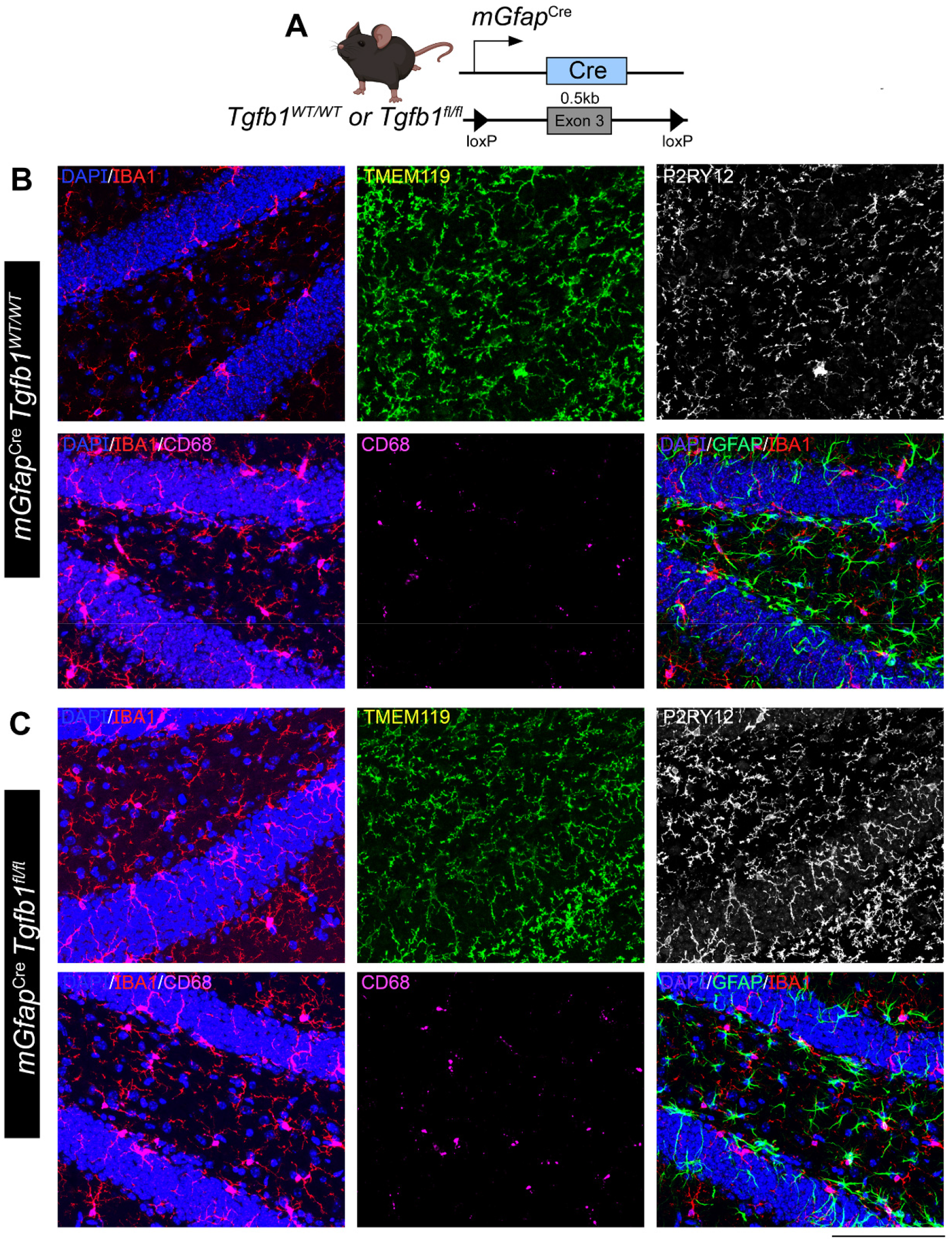
Astrocyte-specific *Tgfb1* gene deletion in the perinatal constitutive mGfap^Cre^ driver line does not affect the homeostasis of microglia or GFAP expression in astrocytes in the adult mouse brain (hippocampus). (A) Astrocyte constitutive KO mouse model used and experimental timeline. (B, C) Representative images of control *mGfap*^Cre^*Tgfb1*^wt/wt^ (B) and cKO *mGfap*^CreER^ *Tgfb1*^fl/fl^ tissue showing IBA1, TMEM119, P2RY12, CD68, and GFAP immunostaining. Representative results from n=3-5 mice/group. Scale bar = 100µm.

**Supplemental Figure 8.**
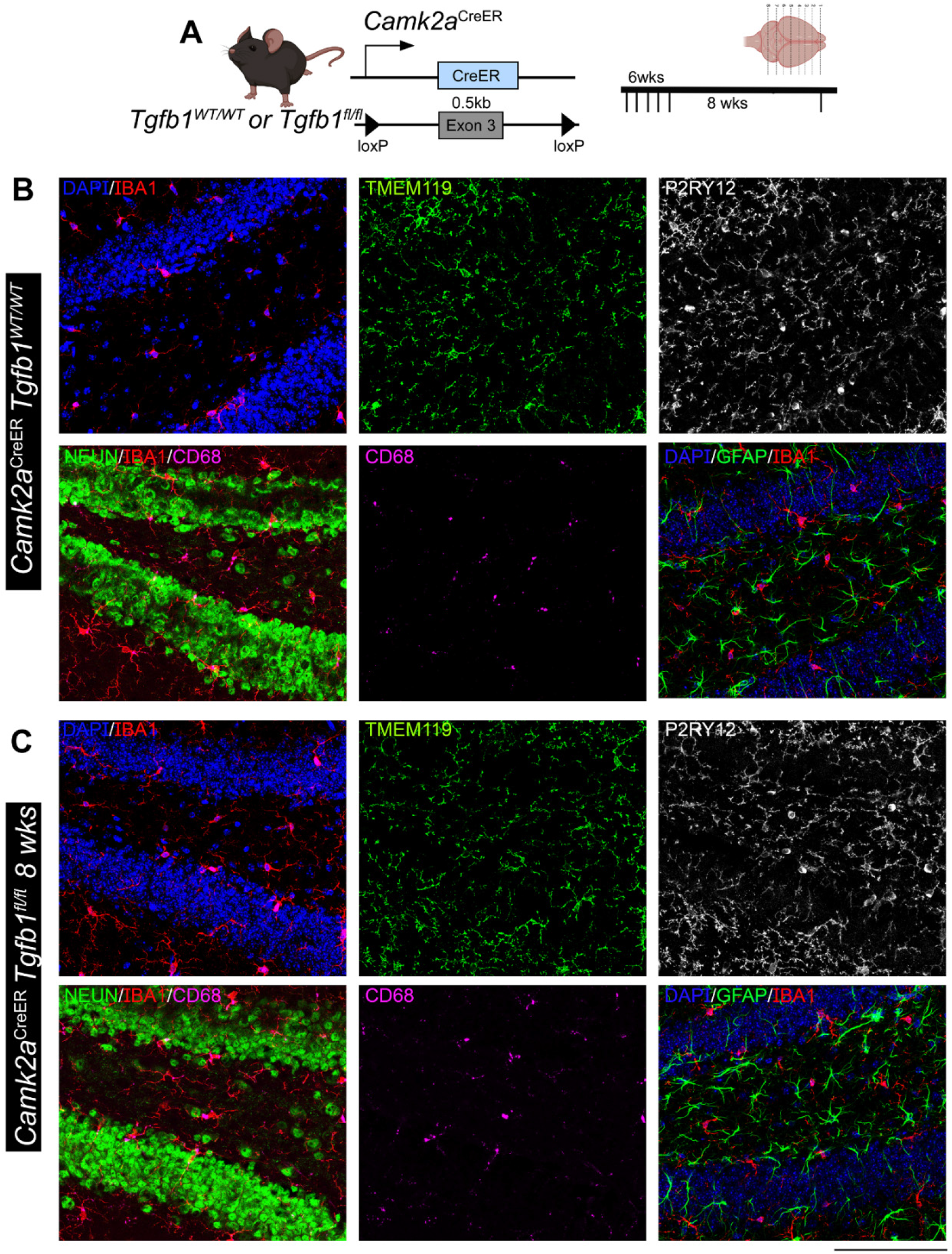
Forebrain neuronal specific *Tgfb1* gene deletion in the *Camk2a*^CreER^ driver does not affect the homeostasis of microglia or GFAP expression in astrocytes in the adult mouse brain (hippocampus). (A) Neuron iKO mouse model used and experimental timeline. (B,C) Representative images of control *Camk2*^Cre^*Tgfb1*^wt/wt^ (B) and iKO *Camk2*^CreER^ *Tgfb1*^fl/fl^ tissue showing IBA1, TMEM119, P2RY12, CD68, and GFAP. Representative results from n=3-5 mice/group. Scale bar = 100µm.

**Supplemental Figure 9.**
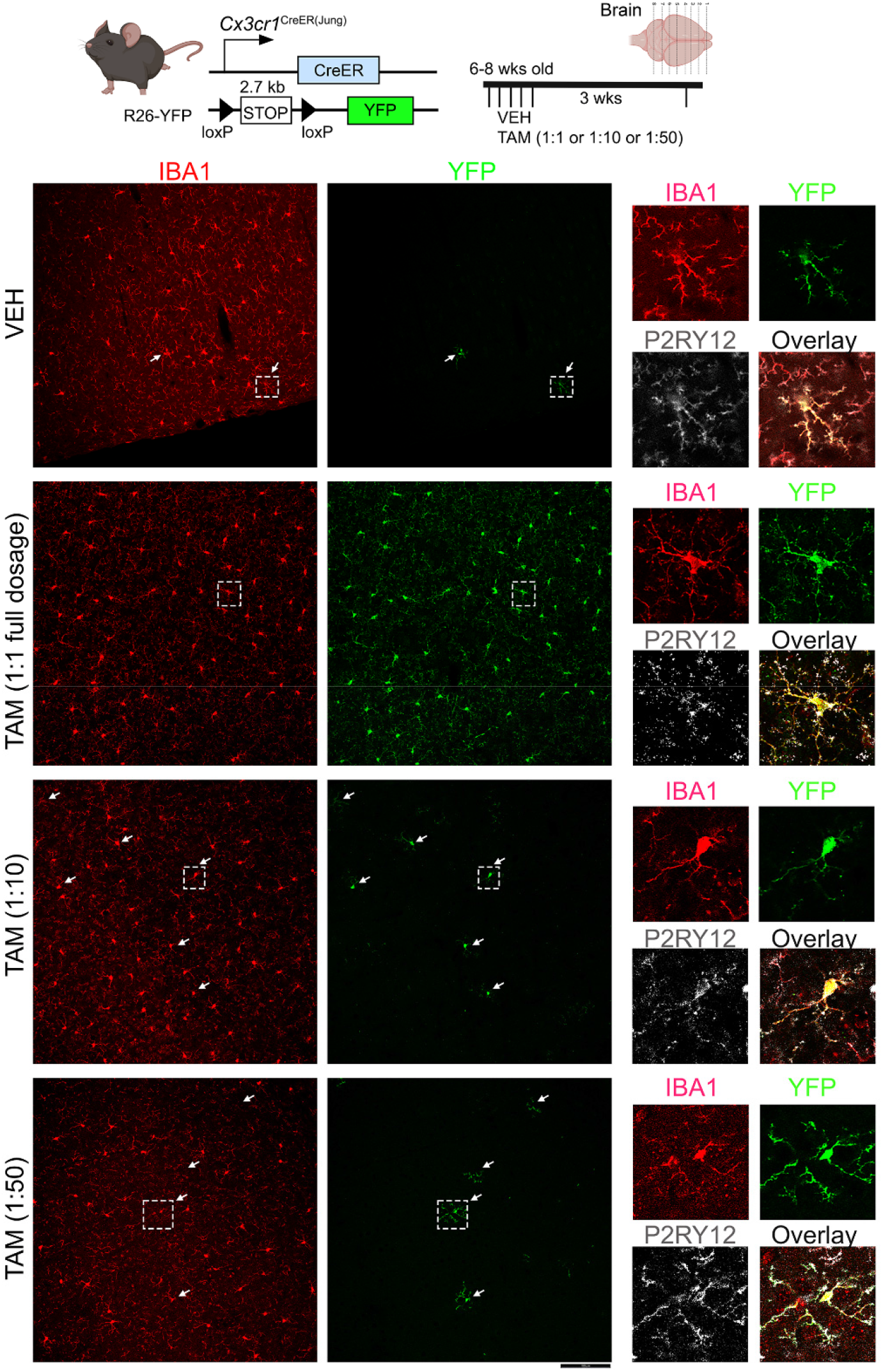
Titration of Tamoxifen dosage to enable sparse recombination of a floxed allele in single microglia in adult mouse brain. (A) The reporter mouse model used to test recombination efficiency in floxed reporter allele. (B-E) Representative IHC images showing IBA1, YFP, and P2RY12 labeling from (B) vehicle-treated, (C) full TAM (180mg/kg) dosage, (D) 1:7-1:10 TAM dosage, and (E) 1:50 TAM dosage treated brain tissue. We observed similar recombination efficiency which lead to sparse labeling in individual microglia in 1:7-1:10 TAM dosage in our study. P2RY12 expression indicates that parenchyma microglia are recombined. Representative results from n=3-5 mice/group. Scale Bar = 100µm.

**Supplemental Figure 10.**
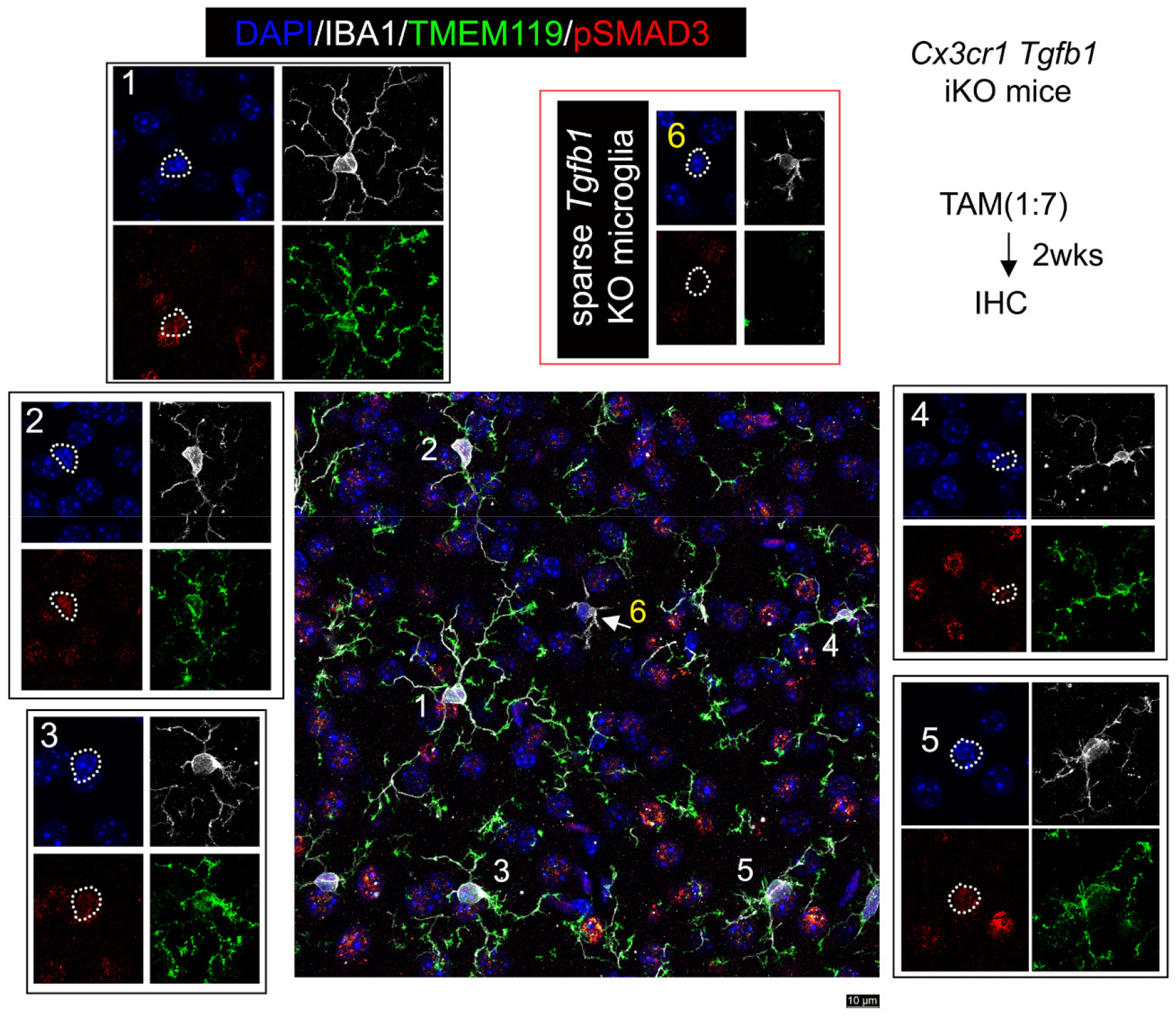
Additional representative pSMAD3 immunostaining labeling confirms the downregulation of TGF-β downstream signaling in dyshomeostatic individual microglia in the sparse *Tgfb1* iKO model. (A) Mouse model used to examine sparse iKO in microglia and experimental timeline with TAM dosage. (B) Representative image showing co-immunohistochemical staining with DAPI, IBA1, TMEM119, and pSMAD3. (B1-5) Surrounding normal microglia showing TMEM119 expression and pSMAD3 immunostaining. (B6) A single microglia cell with loss of TMEM119 expression and loss of pSMAD3 labeling. White arrow (microglia #6) marks the sparse recombined individual iKO microglia. Representative results from n=3 iKO mice. Scale bar = 10µm.

**Supplemental Figure 11.**
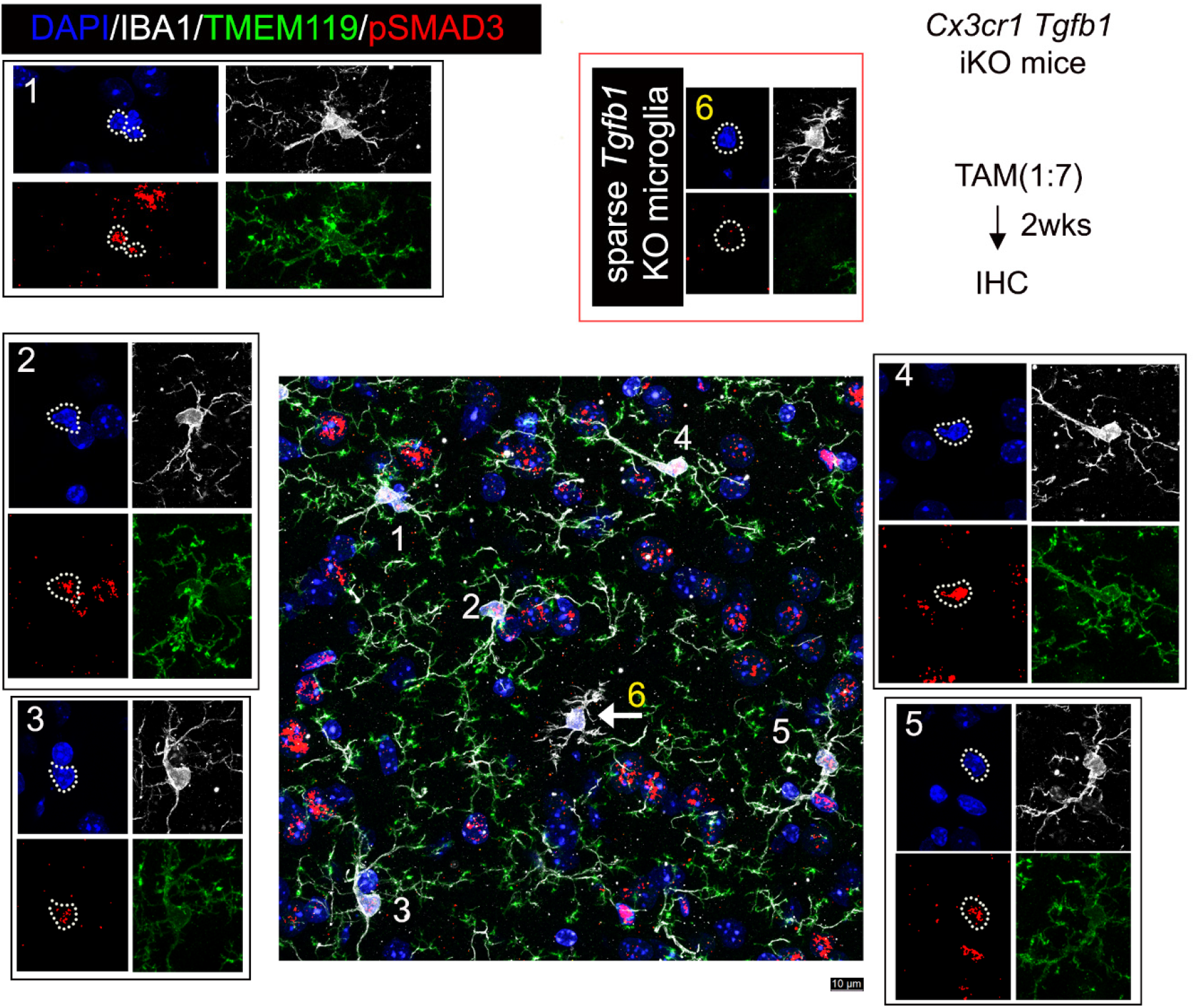
Additional representative pSMAD3 immunostaining labeling confirms the downregulation of TGF-β downstream signaling in dyshomeostatic individual microglia in the sparse *Tgfb1* iKO model. (A) Mouse model used to examine sparse iKO in microglia and experimental timeline with TAM dosage. (B) Representative image showing co-immunohistochemical staining with DAPI, IBA1, TMEM119, and pSMAD3. (B1-5) Surrounding normal microglia showing TMEM119 expression and pSMAD3 immunostaining. (B6) A single microglia cell with loss of TMEM119 expression and loss of pSMAD3 labeling. White arrow (microglia #6) marks the sparse recombined individual iKO microglia. Representative results from n=3 iKO mice. Scale bar = 10µm.

**Supplemental Figure 12.**
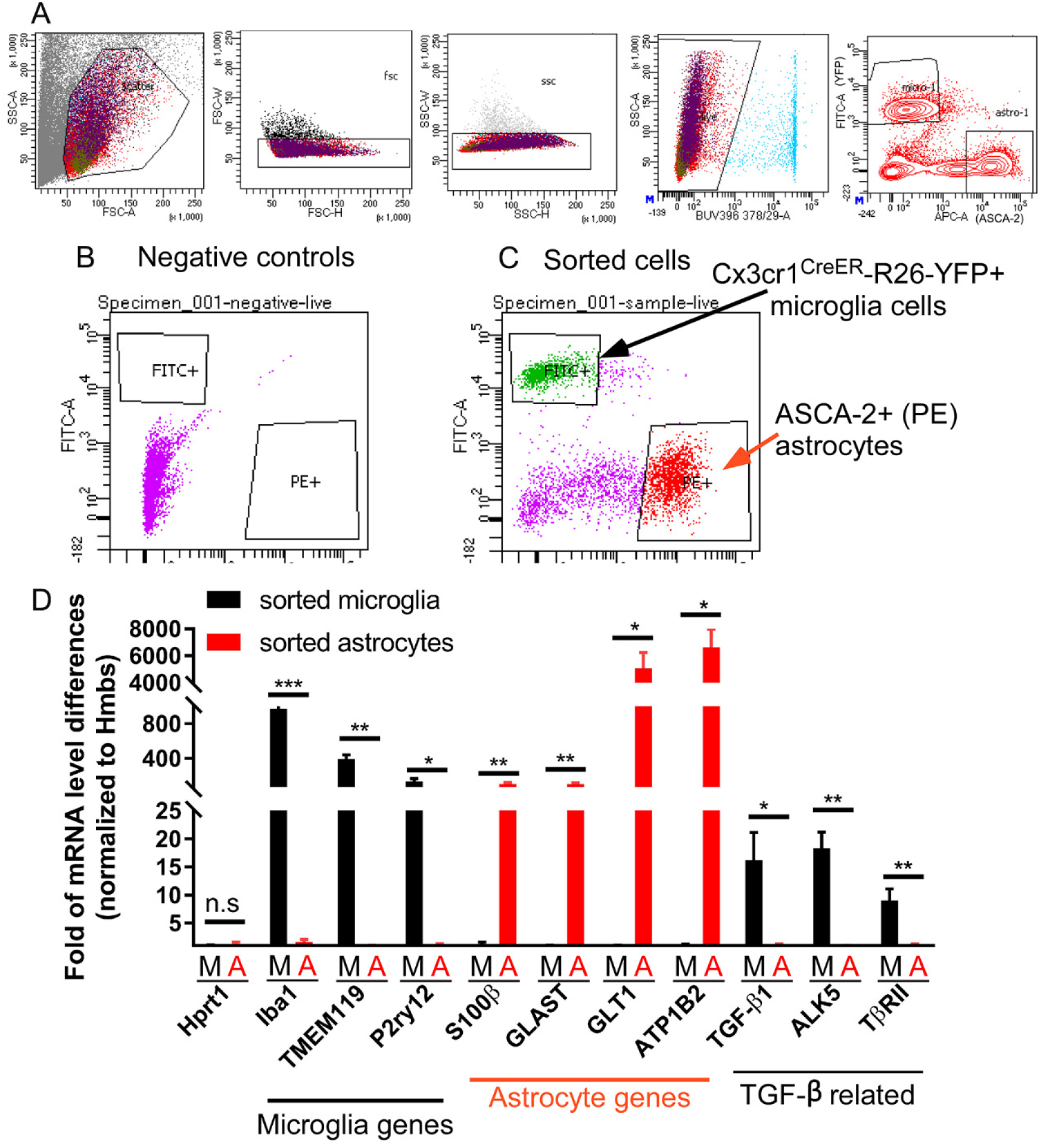
Simultaneous sorting of microglia and astrocyte from the same mouse brain and subsequent RNA extraction and qRT-PCR supporting efficient sorting of microglia and astrocytes from the same brain. (A) Gating strategies for isolation of microglia based on TAM-induced R26-YFP expression and ASCA-2 staining on astrocytes. (B) Examples of unstained samples from a non-TAM treated and non-immunostained brain sample (left) and TAM-treated Cx3cr1CreER-R26-YFP mice that is stained with ASCA-2 antibodies. (C) The purity of sorted cells was validated using qRT-PCR by cell type-specific marker expression for microglia and astrocytes respectively. Note that TGfb1 (ligand) and both type I (ALK5) and type II (TbRII) receptors are substantially enriched in microglia compared to astrocytes. *, **, and *** p< 0.05, 0.01, 0.001. Student’s t-test, n=4 brains.

**Supplemental Figure 13.**
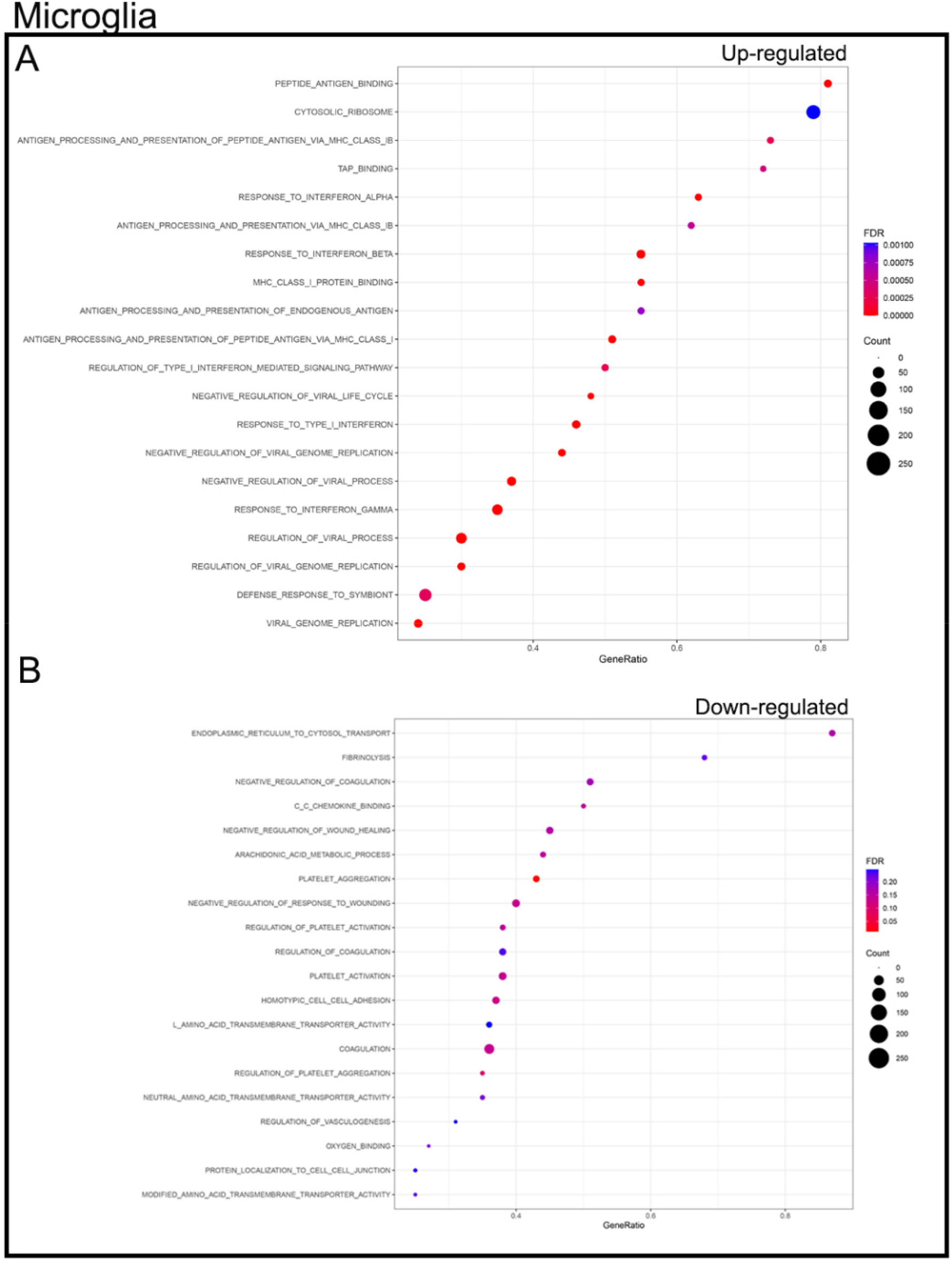
GSEA Analysis of bulk RNA-sequencing data set examining GO pathways in microglia from *Cx3cr1*^CreER(Jung)^*Tgfb1*^fl/fl^ compared to controls. (A) Up-regulated and (B) down-regualted GO pathways in brain microglia three weeks after loss of microglial *Tgfb1*.

**Supplemental Figure 14.**
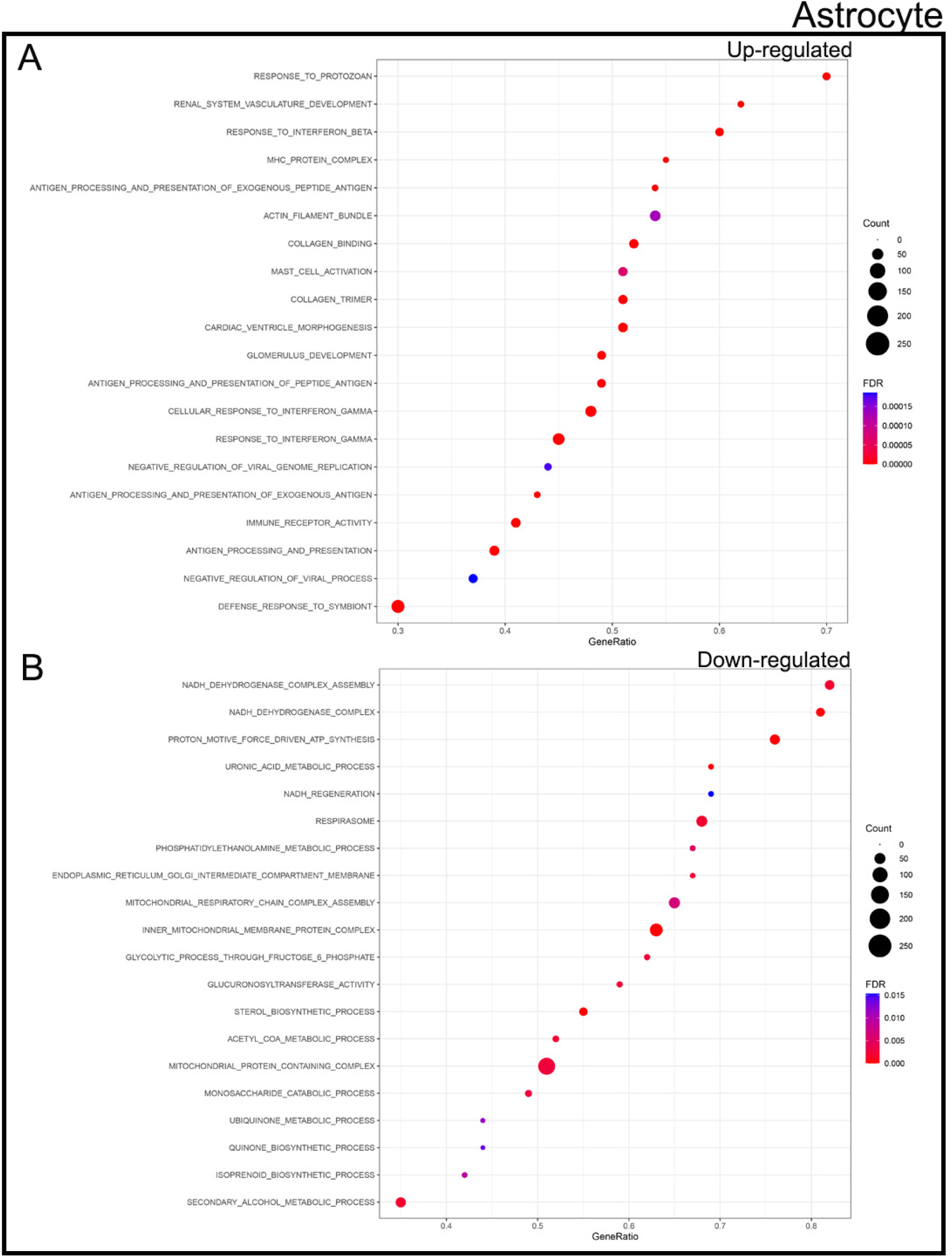
GSEA Analysis of bulk RNA-sequencing data set examining GO pathways in astrocytes from *x3cr1*^CreER(Jung)^*Tgfb1*^fl/fl^ compared to controls. (D) Up- regulated and (E) down-regulated GO pathways of astrocytes from MG-*Tgfb1* iKO mice.

**Supplemental Figure 15.**
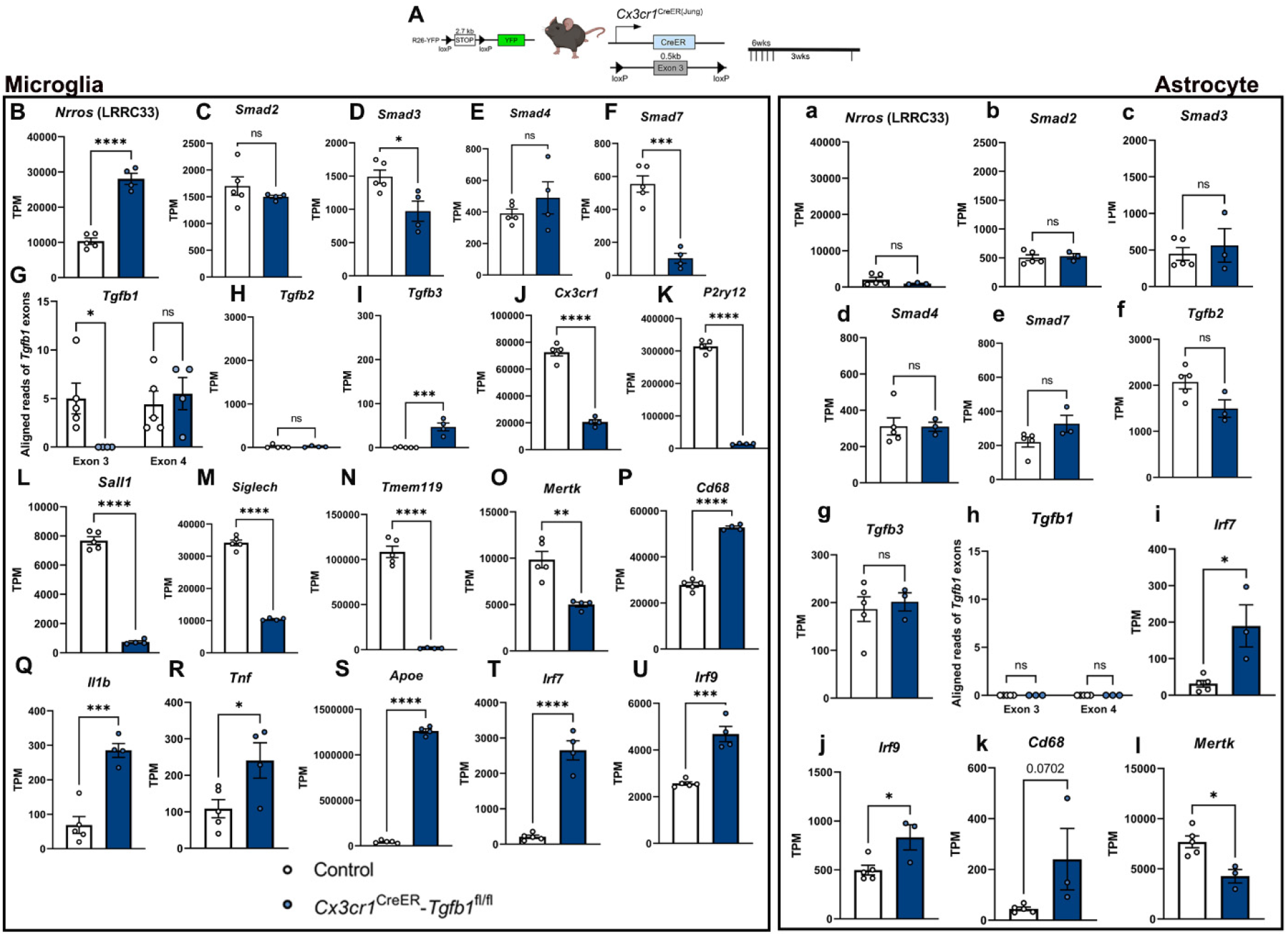
Gene expression changes observed in TGF-β signaling pathway component genes, microglia signature genes, and pro-inflammatory genes. (A) Mouse model used. (B-I) Expression levels of TGF-β signaling components in control microglia and MG- *Tgfb1* iKO microglia. (J-O) Expression levels of microglia signature genes and (P-U) pro-inflammatory genes. (a-h) Astrocytic expression of TGF-β signaling pathway components in control and MG-*Tgfb1* iKO astrocytes. (i-l) Differentially expressed pro-inflammatory genes in astrocytes. *, **, ***, **** p< 0.05, 0.01, 0.001, 0.0001. Student’s t-test, each data point represents a single animal.

## References

1. Paolicelli, R. C. et al. Synaptic pruning by microglia is necessary for normal brain development. Science 333, 1456–1458 (2011).

2. Carroll, J. A., Race, B., Williams, K., Striebel, J. F. & Chesebro, B. Innate immune responses after stimulation with Toll-like receptor agonists in ex vivo microglial cultures and an in vivo model using mice with reduced microglia. J Neuroinflammation 18, 194 (2021).

3. Sierra, A. et al. Microglia shape adult hippocampal neurogenesis through apoptosis-coupled phagocytosis. Cell Stem Cell 7, 483–495 (2010).

4. Nimmerjahn, A., Kirchhoff, F. & Helmchen, F. Resting microglial cells are highly dynamic surveillants of brain parenchyma in vivo. Science 308, 1314–1318 (2005).

5. Hu, X. et al. Microglia/macrophage polarization dynamics reveal novel mechanism of injury expansion after focal cerebral ischemia. Stroke 43, 3063–3070 (2012).

6. Han, Y. et al. TNF- a mediates SDF-1 a – induced NF- k B activation and cytotoxic effects in primary astrocytes Find the latest version: TNF-α mediates SDF-1 α – induced NF-κ B activation and. 108, 425–435 (2001).

7. Lian, H., Litvinchuk, X. A., Chiang, A. C., Aithmitti, N. & Jankowsky, J. L. Astrocyte-Microglia Cross Talk through Complement Activation Modulates Amyloid Pathology in Mouse Models of Alzheimer ’ s Disease. 36, 577–589 (2016).

8. Butovsky, O. et al. Identification of a unique TGF-β-dependent molecular and functional signature in microglia. Nature Neuroscience 17, 131–143 (2014).

9. Zöller, T. et al. Silencing of TGFβ signalling in microglia results in impaired homeostasis.Nature Communications 9, 1–13 (2018).

10. Mattson, M. P. et al. Cellular signaling roles of TGFβ, TNFα and βAPP in brain injury responses and Alzheimer’s disease. Brain Research Reviews 23, 47–61 (1997).

11. Qin, Y. et al. A Milieu Molecule for TGF-B Required for Microglia Function in the Nervous System. Cell 174, 156–157 (2018).

12. Meyers, E. A. & Kessler, J. A. TGF-β family signaling in neural and neuronal differentiation, development, and function. Cold Spring Harbor Perspectives in Biology 9, (2017).

13. Kashima, R. & Hata, A. The role of TGF-β superfamily signaling in neurological disorders. Acta Biochimica et Biophysica Sinica 50, 106–120 (2018).

14. Vivien, D. & Ali, C. Transforming growth factor-β signalling in brain disorders. Cytokine and Growth Factor Reviews 17, 121–128 (2006).

15. Spittau, B., Dokalis, N. & Prinz, M. The Role of TGFβ Signaling in Microglia Maturation and Activation. Trends in Immunology 41, 836–848 (2020).

16. Duan, Z. et al. Specificity of TGF-β1 signal designated by LRRC33 and integrin αVβ8. Nat Commun 13, 4988 (2022).

17. Zhang, Y. et al. An RNA-sequencing transcriptome and splicing database of glia, neurons, and vascular cells of the cerebral cortex. Journal of Neuroscience 34, 11929–11947 (2014).

18. Broad Institute. Study: Synucleinopathy-associated astrocytes. Single Cell Portal https://singlecell.broadinstitute.org/single_cell/study/SCP1879/synucleinopathy-associated-astrocytes (2018).

19. Bedolla, A. et al. Diphtheria toxin induced but not CSF1R inhibitor mediated microglia ablation model leads to the loss of CSF/ventricular spaces in vivo that is independent of cytokine upregulation. J Neuroinflammation 19, 3 (2022).

20. Spiteri, A. G. et al. PLX5622 Reduces Disease Severity in Lethal CNS Infection by Off-Target Inhibition of Peripheral Inflammatory Monocyte Production. Front. Immunol. 13, 851556 (2022).

21. Spangenberg, E. et al. Sustained microglial depletion with CSF1R inhibitor impairs parenchymal plaque development in an Alzheimer’s disease model. Nat Commun 10, 3758 (2019).

22. Dos Santos, S. E., et al. Similar Microglial Cell Densities across Brain Structures and Mammalian Species: Implications for Brain Tissue Function. J. Neurosci. 40, 4622–4643 (2020).

23. Lawson, L. J., Perry, V. H., Dri, P. & Gordon, S. Heterogeneity in the distribution and morphology of microglia in the normal adult mouse brain. Neuroscience 39, 151–170 (1990).

24. Parkhurst, C. N. et al. Microglia promote learning-dependent synapse formation through brain-derived neurotrophic factor. Cell 155, 1596–1609 (2013).

25. Bedolla, A., et al. Finding the right tool: a comprehensive evaluation of microglial inducible cre mouse models. http://biorxiv.org/lookup/doi/10.1101/2023.04.17.536878 (2023) doi:10.1101/2023.04.17.536878.

26. Sahasrabuddhe, V. & Ghosh, H. S. Cx3Cr1-Cre induction leads to microglial activation and IFN-1 signaling caused by DNA damage in early postnatal brain. Cell Reports 38, 110252 (2022).

27. Yona, S. et al. Fate Mapping Reveals Origins and Dynamics of Monocytes and Tissue Macrophages under Homeostasis. Immunity 38, 79–91 (2013).

28. Faust, T. E., et al. A comparative analysis of microglial inducible Cre lines. http://biorxiv.org/lookup/doi/10.1101/2023.01.09.523268 (2023) doi:10.1101/2023.01.09.523268.

29. Diniz, L. P., Matias, I., Siqueira, M., Stipursky, J. & Gomes, F. C. A. Astrocytes and the TGF-β1 Pathway in the Healthy and Diseased Brain: a Double-Edged Sword. Molecular Neurobiology 56, 4653–4679 (2019).

30. Cekanaviciute, E. et al. Astrocytic TGF-β Signaling Limits Inflammation and Reduces Neuronal Damage during Central Nervous System Toxoplasma Infection. The Journal of Immunology 193, 139–149 (2014).

31. Doyle, K. P., Cekanaviciute, E., Mamer, L. E. & Buckwalter, M. S. TGFβ signaling in the brain increases with aging and signals to astrocytes and innate immune cells in the weeks after stroke. Journal of Neuroinflammation 7, 1–13 (2010).

32. Srinivasan, R. et al. New Transgenic Mouse Lines for Selectively Targeting Astrocytes and Studying Calcium Signals in Astrocyte Processes In Situ and In Vivo. Neuron 92, 1181–1195 (2016).

33. Gregorian, C. et al. Pten deletion in adult neural stem/progenitor cells enhances constitutive neurogenesis. J Neurosci 29, 1874–1886 (2009).

34. Hill, S. A. et al. Sonic hedgehog signaling in astrocytes mediates cell type-specific synaptic organization. Elife 8, e45545 (2019).

35. Madisen, L. et al. A robust and high-throughput Cre reporting and characterization system for the whole mouse brain. Nat Neurosci 13, 133–140 (2010).

36. Kaiser, T. & Feng, G. Tmem119-EGFP and Tmem119-CreERT2 Transgenic Mice for Labeling and Manipulating Microglia. eNeuro 6, ENEURO.0448-18.2019 (2019).

37. McKinsey, G. L. et al. A new genetic strategy for targeting microglia in development and disease. eLife 9, 1–34 (2020).

38. Batiuk, M. Y. et al. An immunoaffinity-based method for isolating ultrapure adult astrocytes based on ATP1B2 targeting by the ACSA-2 antibody. Journal of Biological Chemistry 292, 8874–8891 (2017).

39. Butovsky, O. & Weiner, H. L. Microglial signatures and their role in health and disease. Nat Rev Neurosci 19, 622–635 (2018).

40. Deczkowska, A. et al. Disease-Associated Microglia: A Universal Immune Sensor of Neurodegeneration. Cell 173, 1073–1081 (2018).

41. Boche, D. & Gordon, M. N. Diversity of transcriptomic microglial phenotypes in aging and Alzheimer’s disease. Alzheimer’s & Dementia 18, 360–376 (2022).

42. Pettas, S. et al. Profiling Microglia through Single-Cell RNA Sequencing over the Course of Development, Aging, and Disease. Cells 11, 2383 (2022).

43. Carroll, J. A., Race, B., Williams, K., Striebel, J. & Chesebro, B. RNA-seq and network analysis reveal unique glial gene expression signatures during prion infection. Mol Brain 13, 71 (2020).

44. Brionne, T. C., Tesseur, I., Masliah, E. & Wyss-coray, T. Loss of TGF-β 1 Leads to Increased Neuronal Cell Death and Microgliosis in Mouse Brain. 40, 1133–1145 (2003).

45. Buckwalter, M. S. et al. Chronically increased transforming growth factor-β1 strongly inhibits hippocampal neurogenesis in aged mice. American Journal of Pathology 169, 154–164 (2006).

46. Harvey, B. K., Hoffer, B. J. & Wang, Y. Stroke and TGF-β proteins: Glial cell line-derived neurotrophic factor and bone morphogenetic protein. Pharmacology and Therapeutics 105, 113–125 (2005).

47. Luo, J. TGF-β as a Key Modulator of Astrocyte Reactivity: Disease Relevance and Therapeutic Implications. Biomedicines 10, 1206 (2022).

48. Peterson, A. J. & O’Connor, M. B. Lean on Me: Cell-Cell Interactions Release TGF-β for Local Consumption Only. Cell 174, 18–20 (2018).

49. Massague, J. TGF-β SIGNAL TRANSDUCTION. Annu. Rev. Biochem. 67, 753–791 (1998).

50. Huang, T., Schor, S. L. & Hinck, A. P. Biological Activity Differences between TGF-β1 and TGF-β3 Correlate with Differences in the Rigidity and Arrangement of Their Component Monomers. Biochemistry 53, 5737–5749 (2014).

51. Arnold, T. D. et al. Impaired αVβ8 and TGFβ signaling lead to microglial dysmaturation and neuromotor dysfunction. Journal of Experimental Medicine 216, 900–915 (2019).

52. Lee, S. et al. CX3CR1 Deficiency Alters Microglial Activation and Reduces Beta-Amyloid Deposition in Two Alzheimer’s Disease Mouse Models. The American Journal of Pathology 177, 2549–2562 (2010).

53. Gyoneva, S., et al. Cx3cr1- deficient microglia exhibit a premature aging transcriptome. Life Sci. Alliance 2, e201900453 (2019).

54. Hickman, S. E., Allison, E. K., Coleman, U., Kingery-Gallagher, N. D. & El Khoury, J. Heterozygous CX3CR1 Deficiency in Microglia Restores Neuronal β-Amyloid Clearance Pathways and Slows Progression of Alzheimer’s Like-Disease in PS1-APP Mice. Front. Immunol. 10, 2780 (2019).

55. Rogers, J. T. et al. CX3CR1 Deficiency Leads to Impairment of Hippocampal Cognitive Function and Synaptic Plasticity. J. Neurosci. 31, 16241–16250 (2011).

56. Sellner, S. et al. Microglial CX3CR1 promotes adult neurogenesis by inhibiting Sirt 1/p65 signaling independent of CX3CL1. acta neuropathol commun 4, 102 (2016).

57. Chytil, A., Magnuson, M. A., Wright, C. V. E. & Moses, H. L. Conditional inactivation of the TGF-β type II receptor using Cre:Lox. genesis 32, 73–75 (2002).

58. Liddelow, S. A. et al. Neurotoxic reactive astrocytes are induced by activated microglia. Nature 541, 481–487 (2017).

59. Brioschi, S. et al. A Cre-deleter specific for embryo-derived brain macrophages reveals distinct features of microglia and border macrophages. Immunity 56, 1027–1045.e8 (2023).

60. Feil, R., Wagner, J., Metzger, D. & Chambon, P. Regulation of Cre Recombinase Activity by Mutated Estrogen Receptor Ligand-Binding Domains. Biochemical and Biophysical Research Communications 237, 752–757 (1997).

61. Marsh, S. E., et al. Single cell sequencing reveals glial specific responses to tissue processing & enzymatic dissociation in mice and humans. bioRxiv 1–13 (2020) doi:10.1101/2020.12.03.408542.

62. Krueger, F. Trim Galore!

63. Marcel, M. cutadapt.

64. Dobin, A. et al. STAR: ultrafast universal RNA-seq aligner. Bioinformatics 29, 15–21 (2013).

65. Tarasov, A., Vilella, A. J., Cuppen, E., Nijman, I. J. & Prins, P. Sambamba: fast processing of NGS alignment formats. Bioinformatics 31, 2032–2034 (2015).

66. O’Leary, N. A. et al. Reference sequence (RefSeq) database at NCBI: current status, taxonomic expansion, and functional annotation. Nucleic Acids Res 44, D733–D745 (2016).

67. Liao, Y., Smyth, G. K. & Shi, W. The R package Rsubread is easier, faster, cheaper and better for alignment and quantification of RNA sequencing reads. Nucleic Acids Research 47, e47–e47 (2019).

68. Love, M. I., Huber, W. & Anders, S. Moderated estimation of fold change and dispersion for RNA-seq data with DESeq2. Genome Biol 15, 550 (2014).

69. Subramanian, A., et al. Gene set enrichment analysis: A knowledge-based approach for interpreting genome-wide expression profiles. Proc. Natl. Acad. Sci. U.S.A. 102, 15545–15550 (2005).

70. Mootha, V. K. et al. PGC-1α-responsive genes involved in oxidative phosphorylation are coordinately downregulated in human diabetes. Nat Genet 34, 267–273 (2003).

71. Wickham, H. ggplot2: Elegant Graphics for Data Analysis. (Springer International Publishing : Imprint: Springer, 2016). doi:10.1007/978-3-319-24277-4.

